# Complement- and inflammasome-mediated autoinflammation-paroxysmal nocturnal hemoglobinuria

**DOI:** 10.1101/635573

**Authors:** Britta Hoechsmann, Yoshiko Murakami, Makiko Osato, Alexej Knaus, Michi Kawamoto, Norimitsu Inoue, Tetsuya Hirata, Shogo Murata, Markus Anliker, Thomas Eggerman, Marten Jaeger, Ricarda Floettmann, Alexander Hoellein, Sho Murase, Yasutaka Ueda, Jun-ichi Nishimura, Yuzuru Kanakura, Nobuo Kohara, Hubert Schrezenmeier, Peter M. Krawitz, Taroh Kinoshita

## Abstract

Paroxysmal nocturnal hemoglobinuria (PNH) is an acquired hematopoietic stem cell disorder characterized by complement-mediated hemolysis and thrombosis, and bone marrow failure. Affected cells harbor somatic mutation in X-linked *PIGA* gene, essential for the initial step in glycosylphosphatidylinositol (GPI) biosynthesis. Loss of GPI biosynthesis results in defective cell-surface expression of GPI-anchored complement regulators CD59 and DAF. The affected stem cells generate many abnormal blood cells after clonal expansion, which occurs under bone marrow failure. Here, we report the mechanistic basis of a disease entity, autoinflammation-paroxysmal nocturnal hemoglobinuria (AIF-PNH), caused by germline mutation plus somatic loss of *PIGT* on chromosome 20q. A region containing maternally imprinted genes implicated in clonal expansion in 20q-myeloproliferative syndromes was lost together with normal *PIGT* from paternal chromosome 20. Taking these findings together with a lack of bone marrow failure, the mechanisms of clonal expansion in AIF-PNH appear to differ from those in PNH. AIF-PNH is characterized by intravascular hemolysis and recurrent autoinflammation, such as urticaria, arthralgia, fever and aseptic meningitis. Consistent with PIGT’s essential role in synthesized GPI’s attachment to precursor proteins, non-protein-linked free GPIs appeared on the surface of PIGT-defective cells. PIGT-defective THP-1 cells accumulated higher levels of C3 fragments and C5b-9 complexes, and secreted more IL-1β than PIGA-defective cells after activation of the complement alternative pathway. IL-1β secretion was dependent upon C5b-9 complexes, accounting for the effectiveness of the anti-C5 drug eculizumab for both intravascular hemolysis and autoinflammation. These results suggest that free GPIs enhance complement activation and inflammasome-mediated IL-1β secretion.

## Introduction

Paroxysmal nocturnal hemoglobinuria (PNH) is an acquired hematopoietic stem cell (HSC) disorder characterized by complement-mediated hemolysis, thrombosis and bone marrow failure (1, 2). Affected cells harbor a somatic mutation in the *PIGA* gene, essential for the initial step in glycosylphosphatidylinositol (GPI) biosynthesis that occurs in the endoplasmic reticulum (ER)(Figure 1A(a)) (3). Loss of GPI biosynthesis results in the defective expression of GPI-anchored proteins (GPI-APs) including complement inhibitors CD59 and DAF/CD55 (Figure 1A(b)). The affected stem cells generate large numbers of abnormal blood cells after clonal expansion that occurs under bone marrow failure. The affected erythrocytes are defective in complement regulation and destroyed by the membrane attack complex (MAC or C5b-9) upon complement activation (1). Eculizumab, an anti-complement component 5 (C5) monoclonal antibody (mAb), has been used to prevent intravascular hemolysis and thrombosis (4, 5). Eculizumab binds to C5 and inhibits its activation and subsequent generation of C5b-9 complexes.

**Figure 1.**
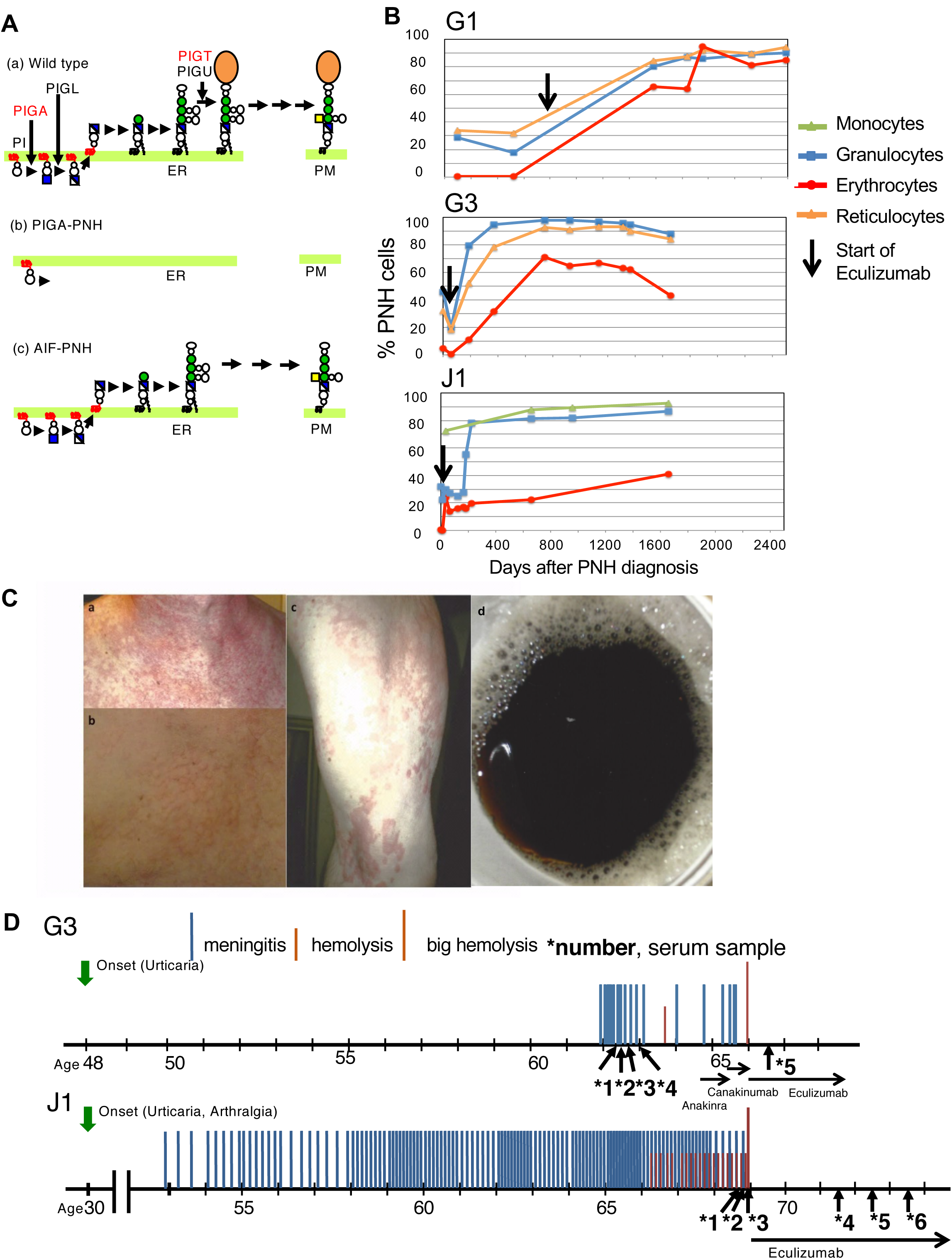
Clinical features of AIF-PNH. **A.** Schematics of normal and defective biosynthesis of GPI anchors. (a) Normal biosynthesis of GPI-APs. GPI is synthesized in the endoplasmic reticulum (ER) from phosphatidylinositol (PI) by sequential reactions and assembled GPI is attached to proteins (orange oval). PIGA acts in the first step of GPI biosynthesis whereas PIGT acts in attachment of GPI to proteins. GPI-APs are transported from the ER to the plasma membrane (PM). (b) No GPI biosynthesis in PNH caused by *PIGA* defect. (c) Accumulation of free GPI in PNH cells caused by *PIGT* defect. GPI is not attached to proteins because of PIGT defect and is transported to the cell surface. **B.** Time course of PNH clone sizes in patients G1, G3 and J1. Percentages of PNH cells in monocytes, granulocytes, erythrocytes and reticulocytes are plotted as a function of time in days. Arrows, start of eculizumab therapy. **C.** Examples of urticaria in patient G3 before the start of the anakinra treatment: (a) and (b), chest; (c), left upper leg. Brightness in (b) was adjusted to more clearly show raised skin in the affected area. (d) hemoglobinuria of patient G3. The pictures were kindly made available by the patient. **D.** Clinical courses of patients G3 in comparison to J1 (Fig. 1 in Kawamoto M et al (8) was modified with additional data) including effective treatments. G3 (top) had meningitis 19 times since 62 years of age to 65 years of age. Eculizumab therapy started at 66 years of age after a severe hemolysis. J1 (bottom) had meningitis 121 times since 53 years of age to 69 years of age when eculizumab therapy started (8). Downward green arrows, onset of urticaria and/or arthralgia; Blue middle height bars, meningitis; Orange short bars, hemolysis; Orange long bars, severe hemolysis; horizontal arrows of various lengths, treatment periods of effective therapies (Anakinra and canakinumab were given with prednisolone); upward arrows with number and asterisk, serum samples taken for cytokine and other protein determination.

Among more than 20 genes involved in GPI biosynthesis and transfer to proteins, *PIGA* is X-linked whereas all others are autosomal (6). Because of X-linkage, one somatic mutation in *PIGA* causes GPI deficiency in both males and females (3). In contrast, two mutations are required for an autosomal gene, but the probability of somatic mutations in both alleles at the same locus is extremely low, which explains why GPI deficiency in most patients with PNH is caused by *PIGA* somatic mutations. Recently, we reported two patients with PNH whose GPI-AP deficiency was caused by germline and somatic mutations in the *PIGT* gene localized on chromosome 20q (7, 8). Both patients had a heterozygous germline loss-of-function mutation in *PIGT*, along with loss of the normal allele of *PIGT* by a deletion of 8 or 18 Mb occurring in HSCs (7, 8). PIGT, forming a GPI transamidase complex with PIGK, PIGS, PIGU and GPAA1, acts in the transfer of preassembled GPI to proteins in the ER (Figure 1A(a)) (9). In *PIGT*-defective cells, GPI is synthesized but is not transferred to precursor proteins, resulting in GPI-AP deficiency on the cell surface (Figure 1A(c)). We showed recently that non-protein-linked, free GPI remaining in the ER of *PIGT*-defective Chinese hamster ovary (CHO) cells is transported to and displayed on the cell surface (Figure 1A(c)) (10).

Two reported PNH patients with PIGT defect suffered from recurrent inflammatory symptoms that are unusual in patients with PNH (7, 8). Here, we report two more patients with PNH who lost *PIGT* function via a similar genetic mechanism, and present insights into the expansion of *PIGT*-defective clones common among four patients. We also present integrated clinical characteristics of these four patients and show that *PIGT*-defective mononuclear leukocytes, but not *PIGA*-defective mononuclear leukocytes, secreted IL1β in response to inflammasome activators. Using a *PIGT*-knockout THP-1 cell model, we show that complement activation is enhanced on the surface of *PIGT*-defective cells leading to MAC-dependent elevated secretion of IL1β. Against this background, we propose a distinct disease entity, autoinflammation-paroxysmal nocturnal hemoglobinuria (AIF-PNH).

## Results

### Case Report

Japanese patient J1 (8) and German patients G1 (7), G2, and G3 were diagnosed with PNH at the ages of 68, 49, 65, and 66, respectively. The changes in PNH clone sizes in J1, G1 and G3 after PNH diagnosis are shown in Figure 1B. They were treated with eculizumab, which effectively prevented intravascular hemolysis. We reported that in G1, direct Coombs test positive erythrocytes appeared after commencement of eculizumab treatment, suggesting extravascular hemolysis (7). Blood cell counts for G1 and G3 are shown in Figure S1A. Before the diagnosis of PNH, J1, G1 and G3 had suffered inflammatory symptoms including urticaria, arthralgia and fever from the ages of 30, 26, and 48, respectively (Table 1). Urticaria in J1 was associated with neutrophil infiltration (8) and that in G3 with a mixed inflammatory infiltrate (Figure 1C). J1 (8) and G3 suffered from recurrent aseptic meningitis characterized by an abundance of neutrophils in cerebrospinal fluid. Following the initiation of eculizumab treatment for hemolysis 3–5 years previously, J1 and G3 had not suffered any episodes of meningitis (Figure 1D). Urticaria and arthralgia were also ameliorated in all three by eculizumab treatment. G2 had severe arteriosclerosis, which might be related to autoinflammation (Table 1), however, whether G2 had autoinflammatory symptoms is unclear and could not be confirmed because the patient passed away.

**Table 1.**
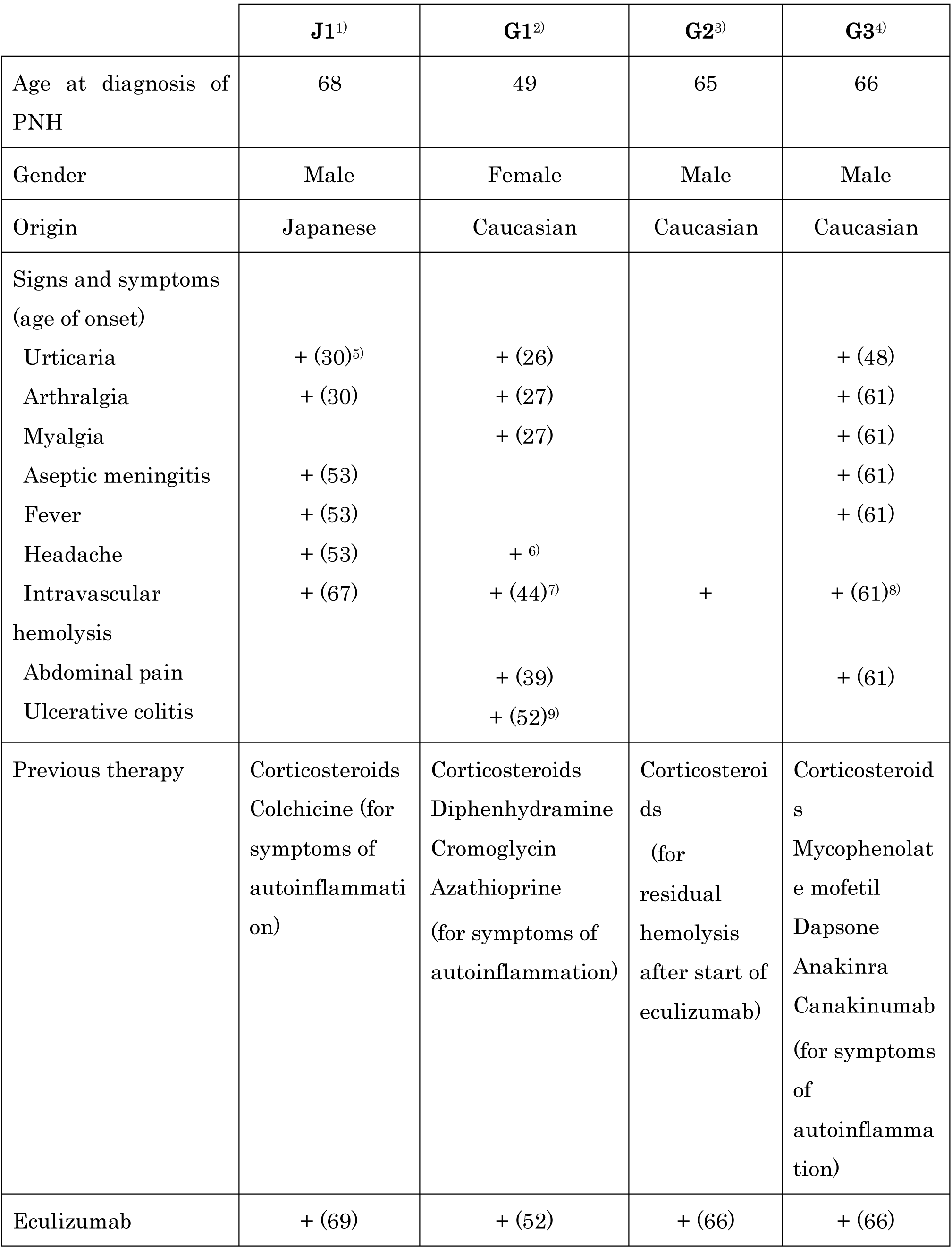
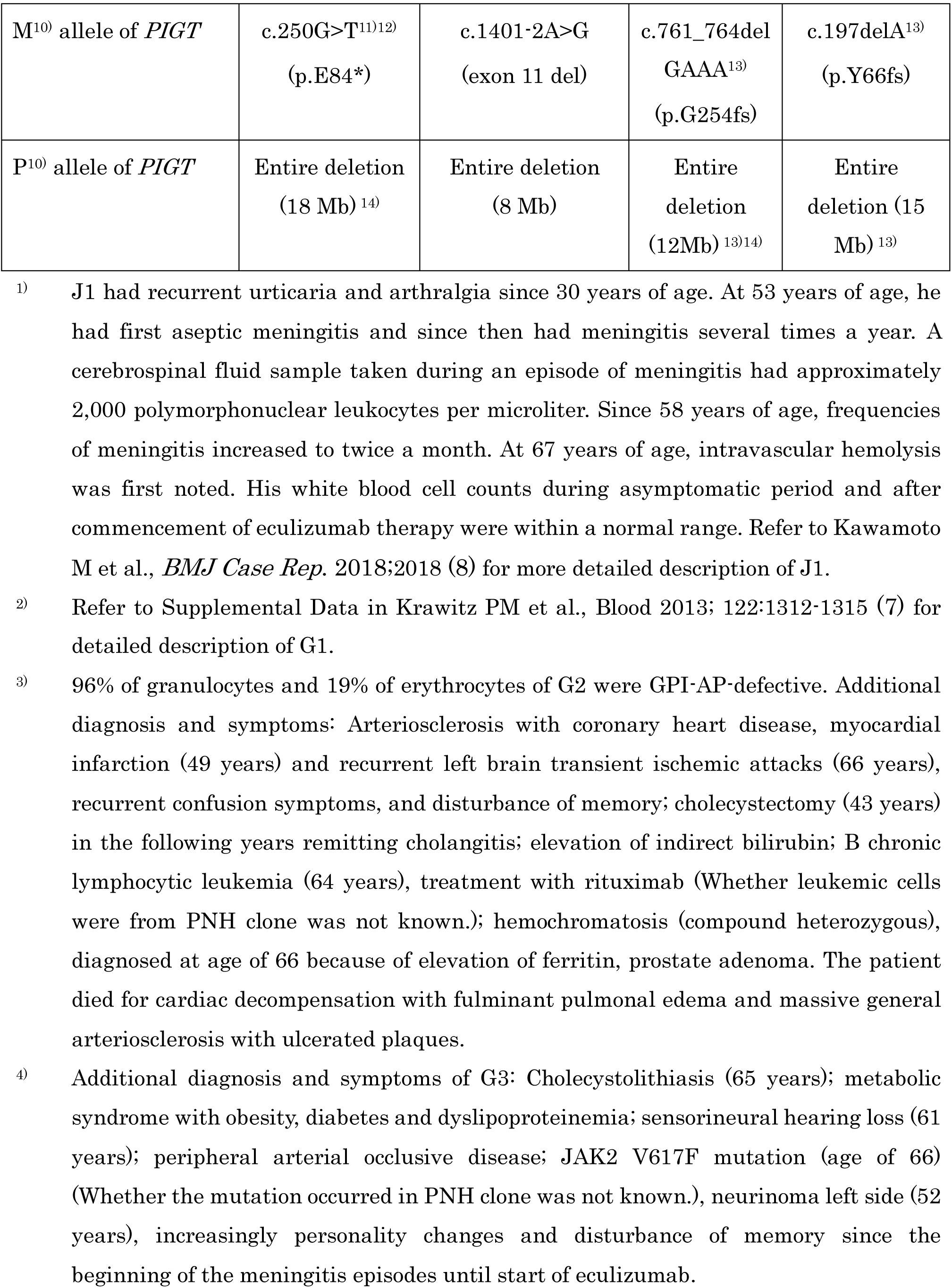
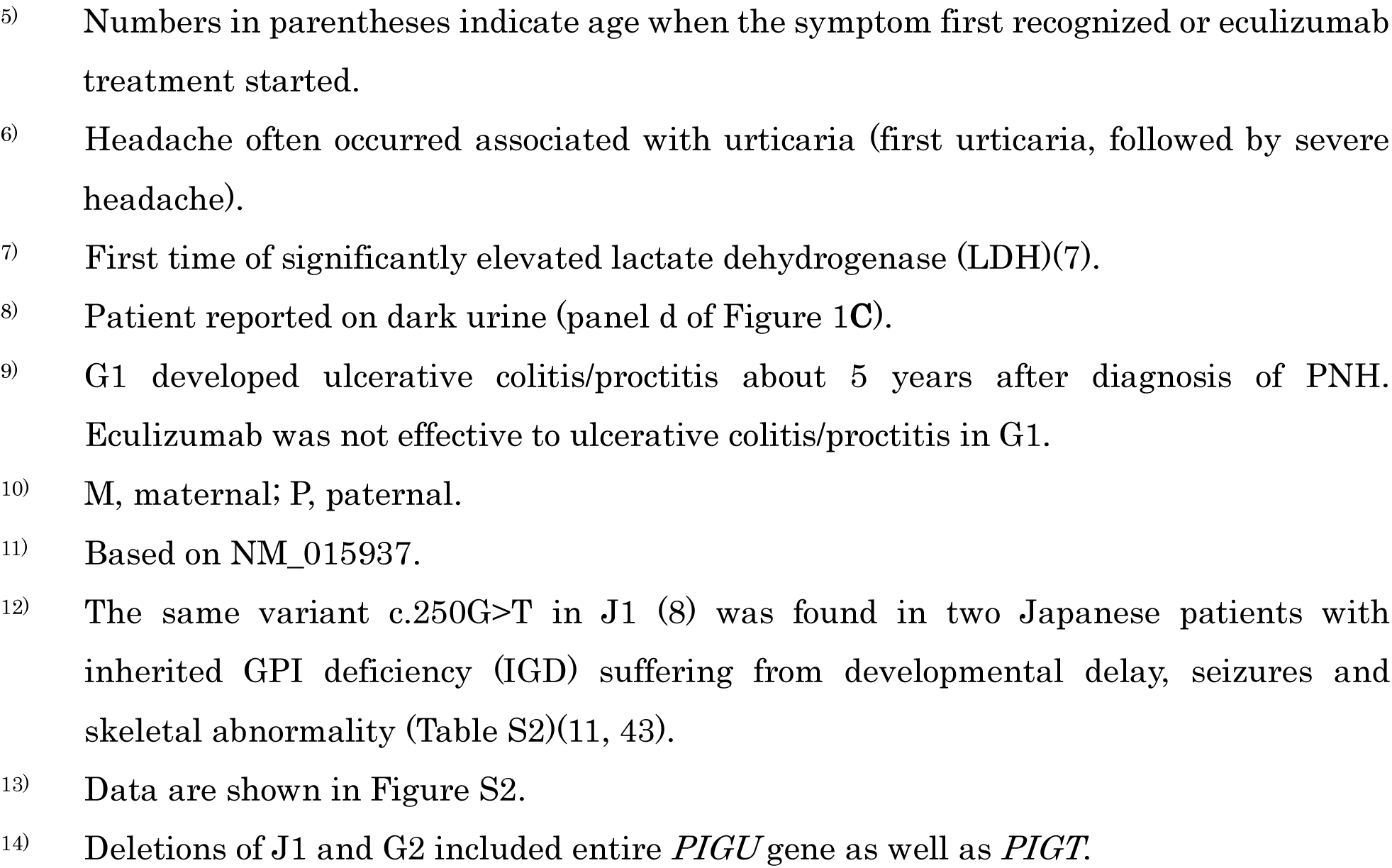
Summary of clinical and genetic findings

### Genetic basis of GPI-AP deficiency

Four patients did not have *PIGA* somatic mutations but had a germline mutation in one allele of *PIGT* located on chromosome 20q: J1, NM_015937 (8): c.250G>T; G1, c.1401-2A>G (7); G2, c.761_764delGAAA; and G3, c.197delA (Figure S2A). These cause E84X, exon 11 skipping, frameshift after G254, and frameshift after Y66, respectively. The functional activities of variant *PIGT* found in J1 and G1 were reported to be very low (7, 11). Variants in G2 and G3 causing frameshifts should also be severely deleterious to PIGT function. In addition to the germline *PIGT* mutation, all four had in the other allele a somatic deletion of 8–18 Mb, which includes the entire *PIGT* gene (Figure S2B) (7, 8). Therefore, in contrast to GPI-AP deficiency caused by a single *PIGA* somatic mutation in PNH, GPI-AP deficiency in all four is caused by a combination of germline loss-of-function *PIGT* mutation and somatic loss of the whole of normal *PIGT* in hematopoietic stem cells (Figure 2A).

**Figure 2.**
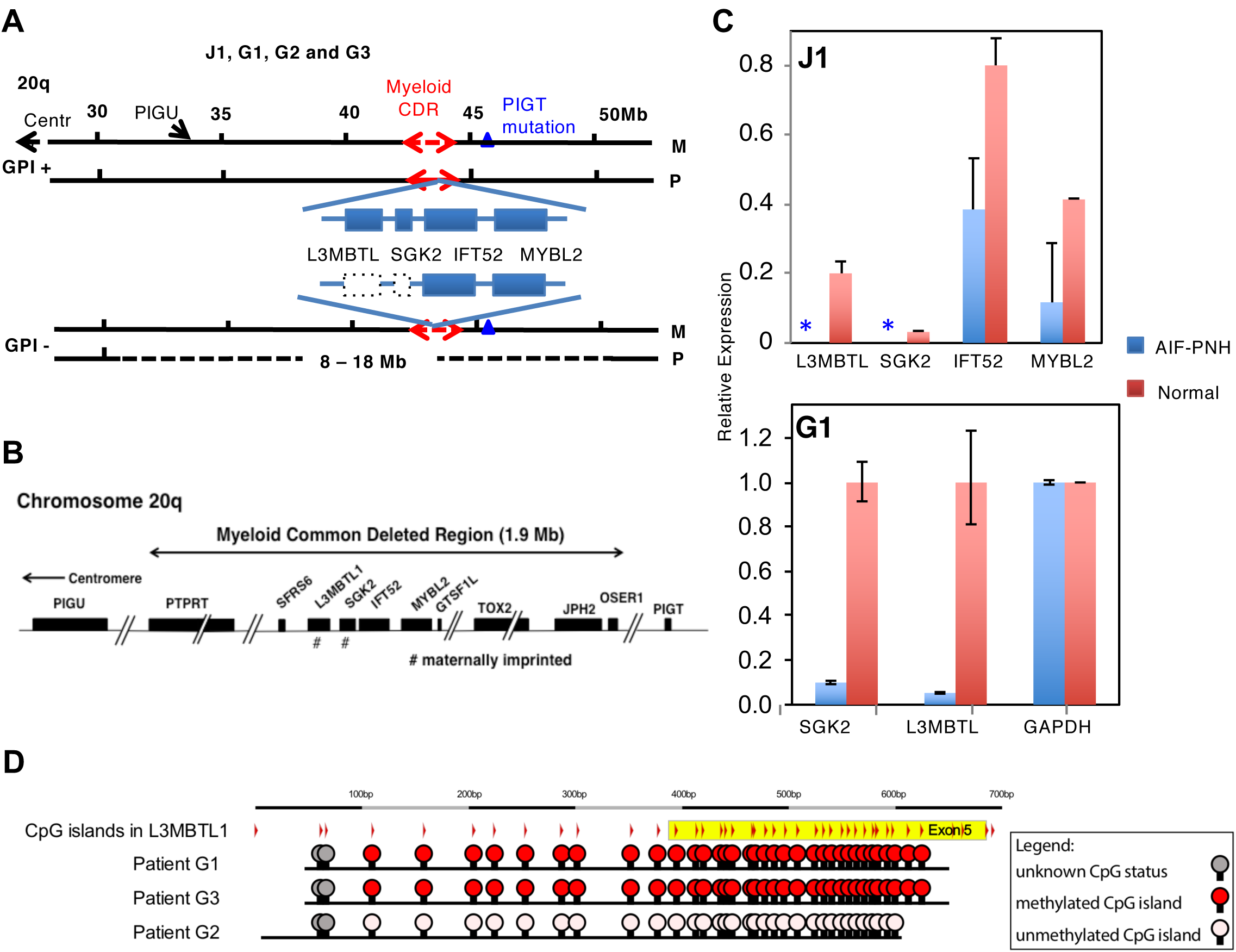
Genetic abnormalities in patients with AIF-PNH. **A.** PIGT mutations in GPI-positive (GPI +) and -defective (GPI -) cells from patients with AIF-PNH. (Top) GPI + cells from patients with AIF-PNH J1, G1, G2 and G3 had a germline *PIGT* mutation (triangle) in the maternal (M) allele. Two maternally imprinted genes, *L3MBTL1* and *SGK2*, within myeloid Common Deleted Region (CDR) are expressed from the paternal (P) allele. Solid and broken red double arrows, P and M alleles of myeloid CDR, respectively. (Bottom) GPI - blood cells from AIF-PNH patients had an 8 Mb to 18 Mb deletion spanning myeloid CDR and *PIGT* and/or *PIGU* in the P chromosome 20q leading to losses of expression of two maternally imprinted *L3MBTL1* and *SGK2* genes (dotted boxes). **B.** Myeloid Common Deleted Region. A 1.9-Mb region in chromosome 20q spanning *PTPRT* gene to *OSER1* gene is termed “myeloid common deleted region (CDR)”. *PIGT* and *PIGU* genes are approximately 1.2 Mb telomeric and 7.4 Mb centromeric to the myeloid CDR, respectively. *L3MBTL1* and *SGK2* genes marked # are maternally imprinted. **C.** qRT-PCR analysis of genes within myeloid CDR in GPI-AP-defective granulocytes from J1 and granulocytes from a healthy control (top) and whole blood cells from G1 and a healthy control (bottom). L3MBTL1 and SGK2, maternally imprinted genes; IFT52 and MYBL2, non-imprinted genes. Relative expression is determined taking means of ABL levels as 1 (J1) or of GAPDH as 1 (G1). Blue bars, cells from J1 and G1; orange bars, cells from healthy individuals; asterisks (*), below detection limits. Mean +/-SD of duplicate (JI) and triplicate (G1) samples in one of the two independent experiments. **D.** Methylation status of the CpG islands in *L3MBTL1* in G1, G2 and G3. Red, methylated CpG islands; pink, unmethylated CpG islands; gray, unknown CpG islands. Bisulfite sequencing data are shown in Figure S3A.

### A possible mechanism of clonal expansion in AIF-PNH

Deletion of chromosome 20q represents the most common chromosomal abnormality associated with myeloproliferative disorders. The deleted region of 8–18 Mb included a “myeloid common deleted region” (CDR) (12) (Figure 2A). A fraction (approximately 10%) of patients with myeloproliferative neoplasm (MPN) such as polycythemia vera commonly have a deletion of 2.7 Mb in chromosome 20q (12). Moreover, a fraction (approximately 4%) of patients with myelodysplastic syndrome (MDS) also have a deletion of 2.6 Mb in this region (12). The CDR region shared by MPN and MDS spanning approximately 1.9 Mb has been called “myeloid CDR” (Figure 2B) and its loss was shown to be causally related to clonal expansion of the affected myeloid cells in these 20q^−^ syndromes (13). In contrast to previous cytogenetic analysis on classical PNH cases that showed no aberrations in 20q (14, 15), one allele of the myeloid CDR was lost in PNH cells of all four patients (Figure S2B) (7, 8).

The tumor suppressor-like gene *L3MBTL1* and the kinase gene *SGK2* located within the myeloid CDR (Figure 2A,B) are expressed only in the paternal allele due to gene imprinting (16). It was shown that losses of active paternal alleles of these two genes had a causal relationship with clonal expansion of these 20q^−^ myeloid cells (13). *L3MBTL1* and *SGK2* transcripts were undetectable in GPI-AP-defective granulocytes from J1 and extremely low in whole blood cells from G1, whereas they were found in granulocytes from healthy individuals (Figure 2C). The transcripts of two unimprinted genes, *IFT52* and *MYBL2*, were detected in both GPI-AP-defective granulocytes from J1 and normal granulocytes (Figure 2C, top). The results therefore indicate that the expression of *L3MBTL1* and *SGK2* is lost in GPI-AP-defective cells in J1 and G1.

The results shown in Figure 2C also indicate that the somatically deleted region in J1 and G1 included active *L3MBTL1* and *SGK2*, so it was in the paternal chromosome. Owing to mRNA from patients G2 and G3 not being available, we determined the methylation status of the *L3MBTL1* gene using DNA from blood leukocytes, among which the large majority of cells were of the PNH phenotype. *L3MBTL1* in G1 and G3 samples was hypermethylated (Figure 2D and Figure S3A), indicating that the myeloid CDR allele remaining in their PNH clones was imprinted. In contrast, the G2 sample was hypomethylated. It was reported that, in some MPN patients with myeloid CDR deletion, the remaining allele was hypomethylated; nevertheless, its transcription was suppressed (13). G2 might be in a similar situation, although it was not possible to draw a definitive conclusion on this by RT-PCR analysis as the patient had passed away. These results indicate that the loss of expressed myeloid CDR allele is associated with clonal expansion of *PIGT*-defective cells similar to 20q− MPN and MDS.

### Appearance of free GPI on the surface of PIGT-defective cells

*PIGA* is required for the first step in GPI biosynthesis (17); therefore, no GPI intermediate is generated in *PIGA*-defective cells (Figure 1A(b)). *PIGT* is involved in the attachment of GPI to proteins. GPI is synthesized in the ER, but is not used as a protein anchor in *PIGT*-defective cells (Figure 1A(c)). We used T5 mAb that recognizes free GPI, but not protein-bound GPI, as a probe to characterize free GPI (Figure 3A: see Methods for epitope and other characteristics of this antibody)(18, 19). Using T5 mAb in western blotting and flow cytometry, we first compared *PIGT*-defective CHO cells with *PIGL*-defective CHO cells, in which an early GPI biosynthetic step is defective, like in *PIGA*-defective cells. T5 mAb revealed a strong band of free GPI at a position corresponding to approximately 10 kDa in lysates of *PIGT*-defective cells but not of *PIGL*-defective cells (Figure 3B). DAF and CD59 were not detected in either mutant cell, confirming that un-GPI-anchored precursor proteins were degraded (Figure 3B) (20). T5 mAb stained the surface of *PIGT*-defective CHO cells but not *PIGL*-defective cells, confirming that free GPI transported to the cell surface is detectable by T5 mAb (Figure 3C).

**Figure 3.**
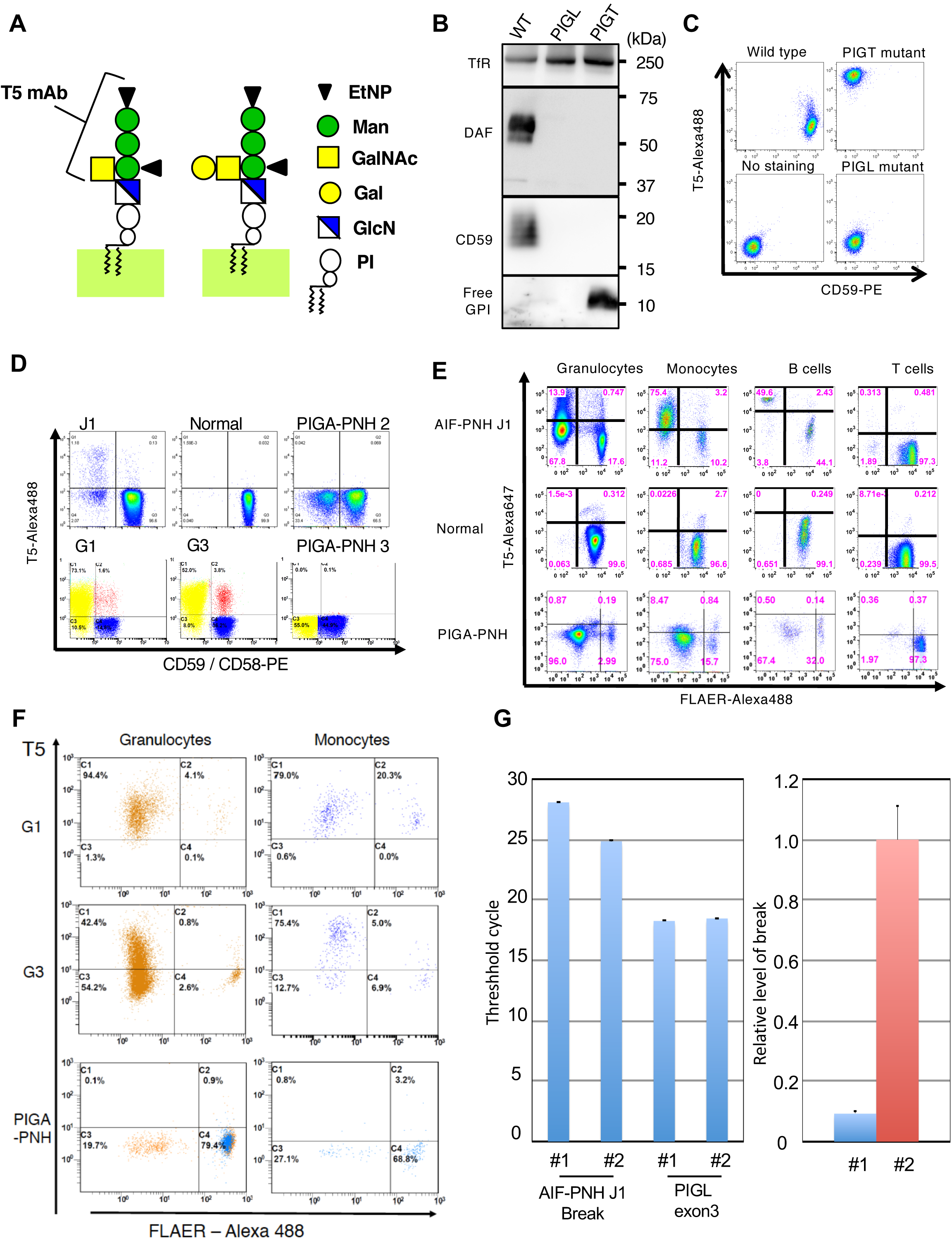
Biochemical abnormalities in PIGT-defective cells. **A.** Schematic representation of binding specificity of T5 mAb. T5 mAb recognizes mammalian free GPI bearing GalNAc-side chain linked to the first mannose (left). T5 mAb does not bind to free GPI when Gal is attached to GalNAc (right). Man, mannose; GlcN, glucosamine; EtNP, ethanolamine phosphate; PI, phosphatidylinositol. **B.** Western blotting analysis of *PIGT*-defective and *PIGL*-defective CHO cells with T5 mAb for free GPI, anti-CD59 and anti-DAF mAbs, and anti-transferrin receptor (TfR) as loading controls. **C.** Flow cytometry of *PIGT*-defective and *PIGL*-defective CHO cells with T5 mAb and anti-CD59 mAb. **D.** Flow cytometry of erythrocytes from J1, G1, G3, a healthy individual and two patients with *PIGA*-PNH with T5 mAb and anti-CD59 (top panels) or anti-CD58 (bottom panels). **E.** Flow cytometry of blood cells from JI, a healthy individual and a patient with *PIGA*-PNH with T5 mAb and FLAER. **F.** Granulocytes and monocytes from G1 and G3, and a patient with *PIGA*-PNH, stained by T5 mAb and FLAER. **G.** Determination of the PNH clone size in J1 by qPCR analysis of the break causing 18 Mbp deletion. (Left) Threshold cycle in PCR for the break and exon 3 of PIGL as a reference. #1, DNA from whole blood leukocytes taken in a stage with autoinflammation only (four months before the onset of recurrent hemolysis); #2, DNA from granulocytes (29% of cells were GPI-AP-defective) taken one month after start of eculizumab therapy. (Right) Relative levels of the break in samples #1 and #2 by setting the level in #2 as 1. Data are shown in mean + SD of triplicate samples in one experiment.

We then analyzed blood cells from J1, G1, and G3, and from patients with *PIGA*-PNH by flow cytometry. All four patients had PNH-type blood cells defective in various GPI-APs (Figures 3D–F and S4A, B). Erythrocytes from J1, G1, and G3 contained 3%, 84%, and 60% PNH cells, respectively, and a sizable fraction of them (36%, 87%, and 87%, respectively) were stained by T5 mAb (Figure 3D). J1 had PNH cells in granulocytes (81%), monocytes (87%), and B-lymphocytes (54%), but not T-lymphocytes (<2%), as revealed by anti-CD59 and GPI-binding probe fluorescence-labeled nonlytic aerolysin (FLAER). Affected monocytes and B-lymphocytes were strongly stained by T5 mAb, whereas affected granulocytes were weakly but clearly stained (Figures 3E and S4B). Normal populations in granulocytes, monocytes and B-lymphocytes were not stained by T5 mAb (Figure 3E). Similar results, showing strong T5 staining of affected monocytes and granulocytes, were obtained with leukocytes from G1 and G3 (Figure 3F). In contrast, PNH cells from *PIGA*-PNH patients and cells from healthy individuals were not positively stained by T5 mAb (Figure 3E, F). Small fractions of wild-type erythrocytes from J1, G1, and G3 (0.13%, 9.7%, and 9.5%, respectively) were positively stained by T5 mAb (Figure 3D). Free GPI might be transferred from PNH cells to wild-type erythrocytes in vivo (21, 22), although the exact mechanism involved needs to be clarified. Thus, the surface expression of the T5 mAb epitope is specific for *PIGT*-defective cells and T5 mAb is useful to diagnose AIF-PNH.

**Figure 4.**
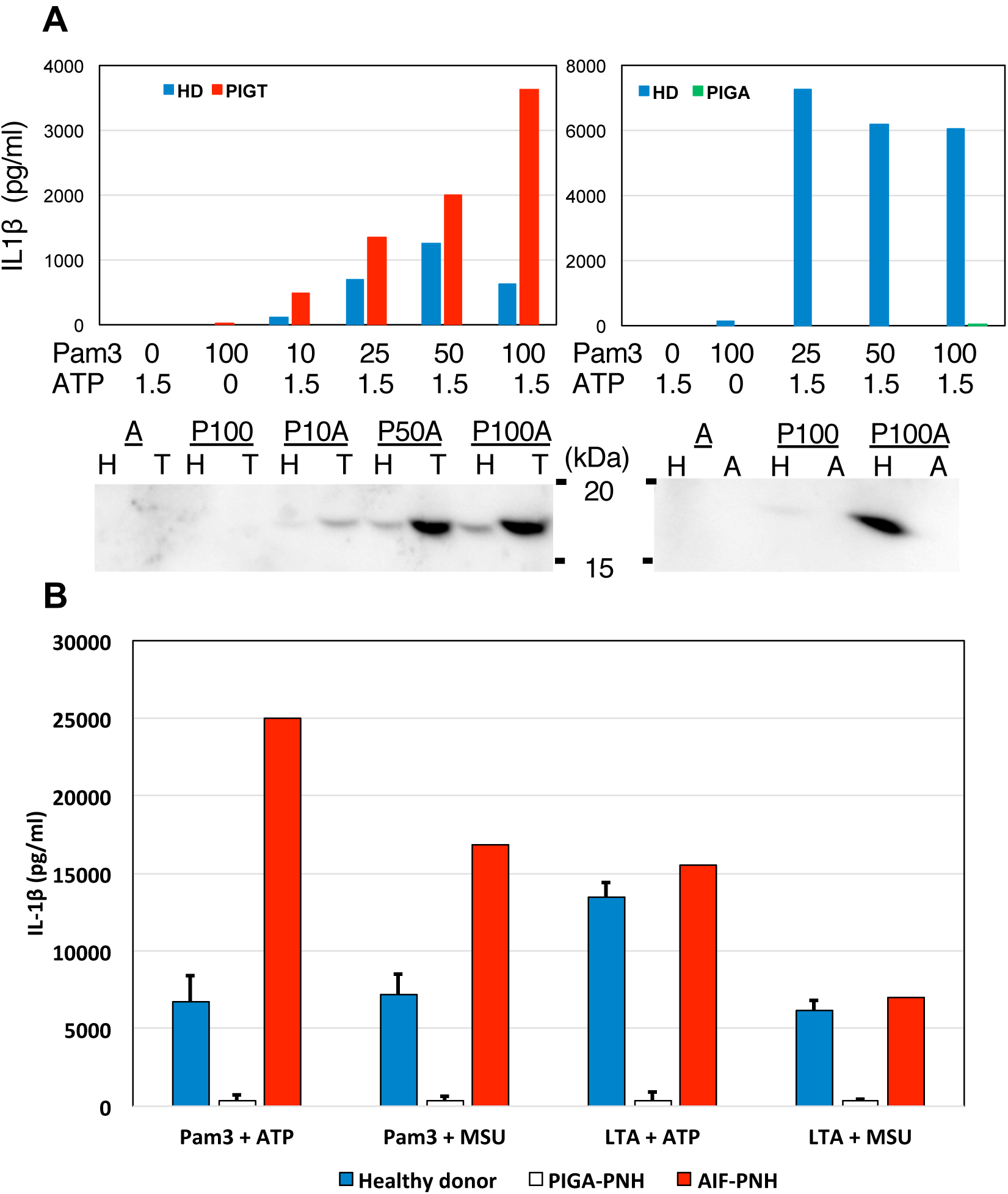
IL1β secretion from AIF-PNH and PIGA-PNH cells. **A.** The peripheral blood mononuclear cells from JI, PIGA-PNH4 and a healthy individual were incubated with 10 to 100 ng/ml Pam_3_CSK4 (Pam3) at 37℃ for 4 hr, and then were incubated with 1.5 mM ATP for 30 min. IL1β in the supernatants was measured by ELISA (top) and western blotting (bottom). (Left) J1 (red bars) and a healthy individual (blue bars). (Right) PIGA-PNH4 (green bars) and a healthy individual (blue bars). <u>A</u>, ATP only; <u>P100</u>, 100 ng/ml Pam_3_CSK4 only; <u>P10A</u>, <u>P50A</u> and <u>P100A</u>, 10, 50 and 100 ng/ml Pam_3_CSK4, respectively and ATP; H, healthy donors; T, AIF-PNH; A, PIGA-PNH. **B.** The peripheral blood mononuclear cells from J1 and two or three patients with *PIGA*-PNH and three healthy controls were stimulated with 200 ng/ml of Pam3 or 1 μg/ml of lipoteichoic acid (LTA) from *Staphylococcus aureus* for 4 hr at 37℃ and after washing were incubated with 3mM ATP or 200 μg/ml monosodium ureate (MSU) for 4 hr at 37℃. IL1β secreted into the medium was determined by ELISA. Data for healthy donors and *PIGA*-PNH were shown as mean + SD. Cells from three patients with *PIGA*-PNH (PIGA-PNH4-6) secreted very low levels of IL1β (403+/-326 pg/ml) after stimulation of NLRP3-inflammasomes with Pam_3_CSK4 and ATP under these strong conditions. In contrast, cells from AIF-PNH J1 secreted IL1β at a high level (25,000 pg/ml) that is higher than levels from three healthy donors (6,693+/-1,711 pg/ml). Similar results were obtained by stimulation with Pam_3_CSK4 and MSU instead of ATP. Moreover, similar results were obtained by stimulation with LTA plus ATP or MSU.

We next investigated whether GPI-AP-defective clone was present in the stage only with autoinflammation. After determining the break points causing the deletion of 18 Mb in J1 (Figure S5), we quantitatively analyzed blood DNA samples for the presence of the break. It was estimated that approximately 3% of total leukocytes obtained 4 months before the onset of recurrent hemolysis had the break, that is, were GPI-AP-defective cells (Figure 3G).

### Inflammasome- and complement-mediated autoinflammation, a feature of AIF-PNH

IL18 levels were elevated in serum samples taken from J1 before and after the commencement of eculizumab therapy (Table 2), suggesting a complement-independent phenomenon. Serum amyloid A was also elevated before eculizumab therapy, but was within the normal range after the commencement of eculizumab therapy (Table 2), suggesting that the elevation was complement-dependent. In G3, increased levels of soluble IL2 receptor and thymidine kinase before, but not after, the start of eculizumab therapy suggested autoinflammation (Table 2). Serum amyloid A (up to 10.5 μg/ml; normal range <5 μg/ml) was also elevated. Combination therapies of prednisolone with anakinra, an IL1 receptor antagonist, or canakinumab, a mAb against IL1β, were effective at reducing urticaria (but not arthralgia and meningitis episodes) of G3. In G1, the IL18 level was above the normal range during eculizumab therapy (195 pg/ml; normal range <150 pg/ml). These lines of evidence suggest that autoinflammatory symptoms are associated with inflammasome activation. We also measured IL18, serum amyloid A, and lactate dehydrogenase (LDH) in serum samples from four patients with *PIGA*-PNH who were not undergoing eculizumab therapy. The levels of IL18 (263–443 pg/ml; normal range <211 pg/ml) and serum amyloid A (5.0–8.3 μg/ml) were within or slightly higher than the normal ranges, whereas LDH levels were markedly elevated (Table 2). These results suggest that autoinflammation is a feature of AIF-PNH.

**Table 2.**
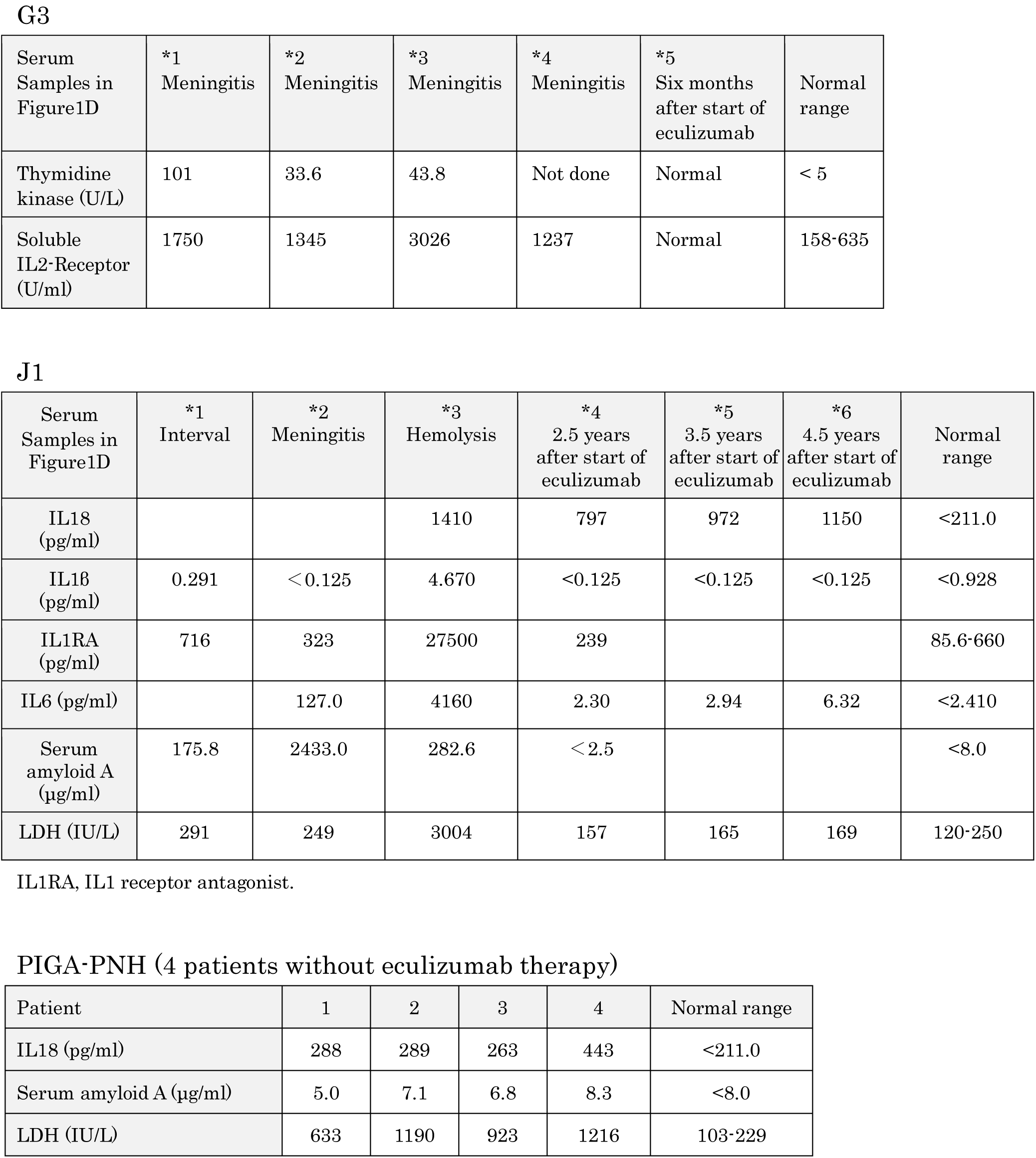
Cytokines and other proteins in serum samples from G3, J1 and PIGA-PNH.

We next compared mononuclear cells from J1, patients with *PIGA*-PNH, and healthy donors for IL1β production upon stimulation by NLRP3-inflammasome activators (23). Cells from three *PIGA*-PNH patients secreted only very low levels of IL1β after stimulation by Pam_3_CSK4 (TLR2 ligand) and ATP or monosodium urate (MSU) (Figure 4A right and 4B). In contrast, cells from J1 secreted 45–60 times as much IL1β and the levels were even higher than those from healthy control cells (Figure 4A left and 4B). A similar difference between PIGT- and PIGA-defective cells was seen upon stimulation by lipoteichoic acid (LTA: another TLR2 ligand) and ATP or MSU (Figure 4B). Low IL1β response of PIGA-defective cells was predicted because they lack CD14, a GPI-anchored co-receptor of TLRs. However, PIGT-defective cells also lacking CD14 showed a strong IL1β response. These results indicate that NLRP3 inflammasomes are easily activated and support the idea that the presence of non-protein-linked free GPI is associated with efficient activation of NLRP3 inflammasomes, contributing to autoinflammatory symptoms in AIF-PNH.

To investigate the roles of complement in inflammasome activation in AIF-PNH, we switched to a model cell system because patients’ blood cells were easily damaged in vitro under conditions of complement activation. PIGTKO and PIGAKO cells were generated from human monocytic THP-1 cells (Figure S6A) and were differentiated to macrophages. They showed IL1β response comparable to those of authentic inflammasome activators (Figure S6B). To analyze the inflammasome response to activated complement, these THP-1-derived macrophages were stimulated with acidified serum (AS), which causes activation of the alternative complement pathway. PIGTKO and PIGAKO cells but not WT cells secreted IL1β (1221.8±91.6, 568.2±101 and 23.7±2.2 pg/ml for PIGTKO, PIGAKO, and WT cells, respectively) (Figure 5A). This result is consistent with impaired complement regulatory activities on PIGTKO and PIGAKO cells, and normal complement regulatory activity on WT cells. PIGTKO cells secreted approximately twice as much IL1β as PIGAKO cells (*p*<0.01). However, IL1β production returned to near the WT cell level after the transfection of *PIGT* and *PIGA* cDNAs into PIGTKO and PIGAKO cells, respectively (Figure 5B). The levels of IL1β mRNA and protein were comparable in WT, PIGTKO, and PIGAKO cells (Figure S7A and S7B). Therefore, PIGT KO enhanced the secretion but not the generation of IL1β.

**Figure 5.**
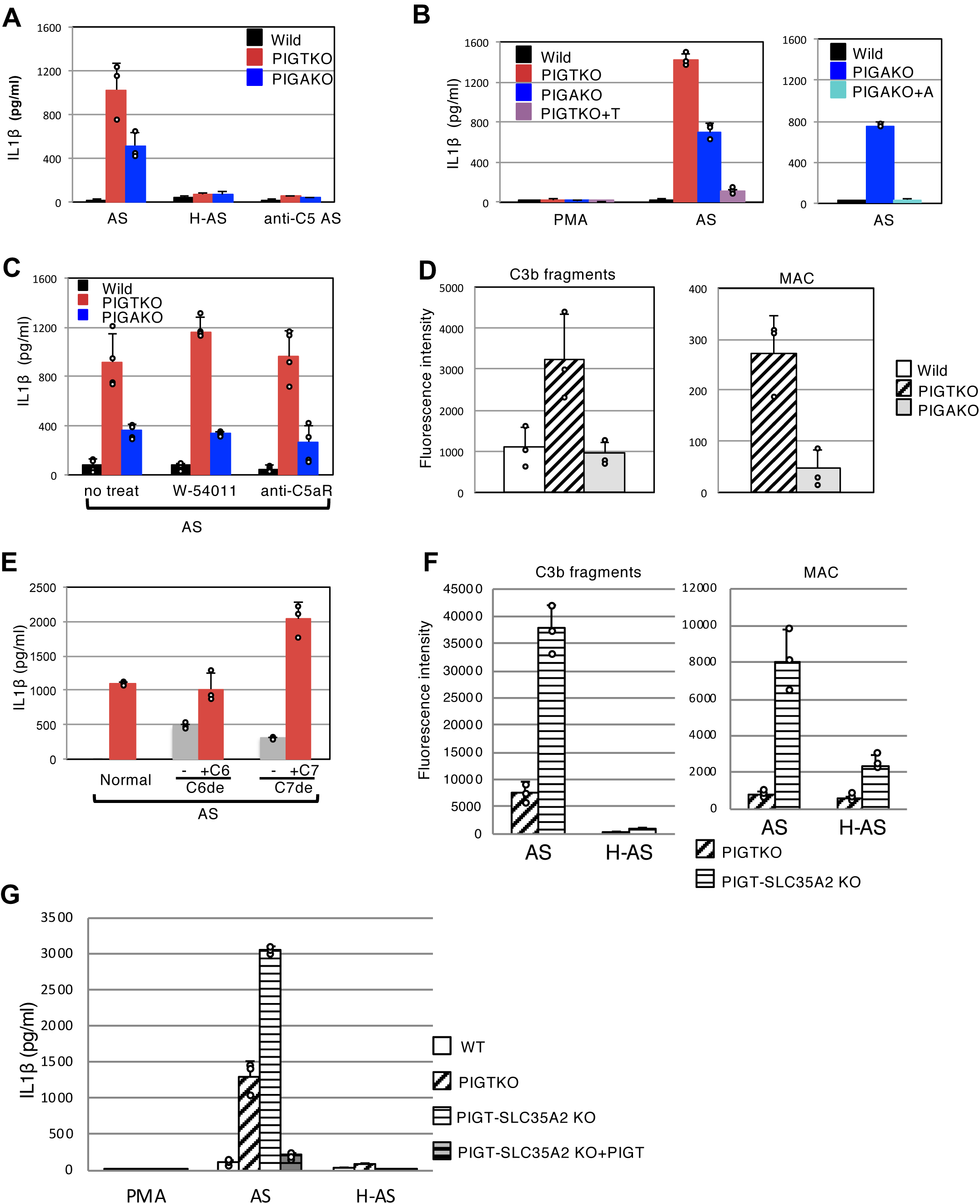
IL1β secretion from PIGT- and PIGA-defective THP-1 cells. **A.** Complement-mediated IL1β secretion from THP-1-derived macrophages. WT, PIGTKO and PIGAKO cells were incubated with acidified serum (AS), heat-inactivated AS (H-AS), or AS containing anti-C5 mAb. Supernatant samples were collected after 5-hr incubation and analyzed for IL1β by ELISA. Mean + SD of three independent experiments. **B.** Reductions of IL1β secretion by transfection of *PIGT* and *PIGA* cDNAs into PIGTKO and PIGAKO cells (PIGTKO+T and PIGAKO+A, respectively). Cells differentiated by PMA were either left untreated (PMA) or incubated with AS (AS) under similar conditions as described in **A** and supernatants analyzed for IL1β. Mean + SD of three independent experiments. **C.** Effect of inhibiting C5aR on IL1β secretion from THP-1-derived macrophages. Cells were incubated with AS alone (no treat), or AS containing C5aR antagonist (W-54011) or anti-C5aR mAb. Supernatant was collected after 5 hr and analyzed for IL1β by ELISA. Mean + SD of duplicate samples from two independent experiments. **D.** Detection of C3b fragments (left) and MAC (right) by flow cytometry on PMA-differentiated THP-1 macrophages after incubation with AS. Geometric mean fluorescence intensity of medium-treated cells was subtracted from that of AS-treated cells. Mean + SD of three independent experiments. **E.** IL1β secretion from PIGTKO THP-1 macrophages stimulated with C6- or C7-depleted AS. PIGTKO THP-1 macrophages were incubated with AS, C6-depleted AS (-/C6de), C6de restored by C6 (+C6/C6de), C7-depleted AS (-/C7de), or C7de restored by C7 (+C7/C7de). Supernatant was collected after overnight incubation. Mean + SD of triplicate samples from two independent experiments (normal and C6-depleted sera) and one experiment (C7-depleted serum). **F.** Binding of C3b fragments (left) and MAC (right) on PIGTKO and PIGT-SLC35A2 double KO THP-1 macrophages after AS treatments. **G.** IL1β secretion from PIGTKO and PIGT-SLC35A2 double KO THP-1 macrophages after AS treatments.

Heat inactivation of complement and the addition of anti-C5 mAb to AS almost completely inhibited IL1β secretion (Figure 5A). These results indicate that IL1β secretion requires the activation of C5 on PIGTKO and PIGAKO cells. The activation of C5 leads to two biologically active products, C5a and MAC (24). To address which of these is important for IL1β secretion, cells were treated with the C5aR antagonist W-54011 (25) or anti-C5aR mAb to inhibit the signal transduction through C5aR. WT, PIGTKO, and PIGAKO cells expressed C5aR at similar levels (Figure S7C). The two methods of functional inhibition of C5aR had little effect on IL1β secretion, indicating that the signal through C5aR plays no major role in this cell system (Figure 5C). Next, AS-treated cells were analyzed for surface binding of C3b fragments and MAC. Exposure to AS resulted in the higher binding of C3b fragments and MAC on PIGTKO cells compared with that on PIGAKO cells (Figures 5D and S8A, B). The level of MAC was several times higher on PIGTKO cells than on PIGAKO cells, suggesting that complement activation was enhanced, leading to the enhanced formation of MAC on PIGTKO cells. To confirm the role of MAC in IL1β secretion, PIGTKO cells were treated with acidified C6- and C7-depleted sera, in which C5a generation is intact whereas MAC formation is impaired. IL1β secretion was greatly reduced by C6 or C7 depletion and was restored by the replenishment of C6 or C7 (Figure 5E). These results suggest that MAC but not C5a plays a critical role in the secretion of IL1β. It is also suggested that free GPI plays some role in complement activation, leading to the enhanced binding of C3b fragments and MAC formation.

Finally, to determine whether the structure of free GPI (presence or absence of Gal capping) affects complement activation and subsequent IL1β secretion, we knocked out SLC35A2 in PIGTKO THP-1 cells. PIGT-SLC35A2 double-KO THP-1 cells were strongly stained by T5 mAb as expected (Figure S6C). The binding of both C3b fragments and MAC increased approximately five times after SLC35A2 KO (Figure 5F). Concomitantly, the secretion of IL1β more than doubled (Figure 5G). These results indicate that the structure of free GPI influenced complement activation efficiency and subsequent IL1β secretion.

## Discussion

We studied patients with PNH caused by *PIGT* mutations and propose that they represent a new disease entity, AIF-PNH. AIF-PNH caused by *PIGT* mutation is distinct from PNH in four regards. First, GPI-AP deficiency in PNH is caused by somatic mutation of the X-linked *PIGA* gene in hematopoietic stem cells, whereas GPI-AP deficiency in AIF-PNH is caused by a germline heterozygous mutation in the *PIGT* gene on chromosome 20q in combination with somatic deletion of the normal *PIGT* gene in hematopoietic stem cells.

Second, *PIGA* mutations cause a defect in the initial step in GPI biosynthesis, whereas *PIGT* mutations cause a defect in the transfer of preassembled GPI to proteins. Therefore, free GPI remains in *PIGT*-defective cells, but not in *PIGA*-defective ones.

Third, the expansion of *PIGA*-defective clones in PNH is often caused by selective survival under autoimmune bone marrow failure with or without the acquisition of benign tumor characteristics by additional somatic mutations (26–29). In contrast, none of the patients with AIF-PNH had documented bone marrow failure (Figure S1A) (7, 8). In addition, the myeloid CDR is lost in *PIGT*-defective clones in AIF-PNH, similar to the case in clonal cells in myeloproliferative 20q− syndromes. The causal relationship between the myeloid CDR loss in AIF-PNH and clonal expansion needs to be proven, particularly because boosted lineages under L3MBTL1 and SGK2 loss differ between in vitro study (13) and AIF-PNH patients. Nevertheless, this unique deletion occurs in AIF-PNH but not in PIGA-PNH. Taking these findings together, it is likely that the mechanism of clonal expansion for AIF-PNH is distinct from that for *PIGA*-PNH cells (see models in Figure S3B).

Fourth, whereas AIF-PNH shares intravascular hemolysis and thrombosis with PNH, AIF-PNH is characterized by autoinflammatory symptoms including recurrent urticaria, arthralgia and aseptic meningitis. AIF-PNH first manifested with autoinflammatory symptoms alone and symptoms of PNH became apparent many years later. It is possible that different clinical symptoms appear depending on the size of the *PIGT*-defective clone. When the clone size is small, autoinflammation but not PNH may occur and, when the clone size becomes sufficiently large, PNH may become apparent. The idea that the *PIGT*-defective clone is small when only autoinflammation is seen was supported by analyzing J1 DNA obtained before the start of recurrent hemolysis, only approximately 3% of total leukocytes being *PIGT*-defective (Figure 3G).

C5 activation must be involved in the autoinflammatory symptoms in AIF-PNH because they were suppressed by eculizumab. It is important to consider GPI-AP deficiency for patients with recurrent autoinflammatory symptoms such as aseptic meningitis even when PNH symptoms are absent because eculizumab may be effective for such cases. Because DAF and CD59 are missing on *PIGT*-defective monocytes, C5a and MAC might be generated once complement activation has been initiated. It was reported that subarachnoidal application of C5a in rabbits and rats induced acute experimental meningitis (30). Various types of myeloid cells are present in the central nervous system (reviewed in (31)). If C5 activation occurs on some of those cells lacking complement regulatory function and C5a is generated, aseptic meningitis might ensue.

The involvement of complement in inflammasome activation has been shown in various blood cell systems (32–35). Indeed, in the THP-1 cell model system, IL1β secretion was induced by complement in both PIGTKO and PIGAKO cells and more strongly in PIGTKO cells, mainly through MAC formation. It appeared that complement activation is enhanced in PIGTKO cells, although the mechanism involved in this is unclear. AIF-PNH mononuclear cells were activated by conventional stimulators of inflammasomes similar to or even stronger than healthy control cells. Because blood mononuclear cells were easily lysed by acidified serum, the effect of complement on inflammasome activation in mononuclear cells could not be addressed. Taking the obtained findings together with the results for THP-1 cells, we speculate that *PIGT*-deficient monocytes show an enhanced inflammasome response when complement is activated. How free GPI is involved in inflammasome and complement activation needs to be clarified to fully understand the mechanistic basis of AIF-PNH.

Inflammatory symptoms, recurrent urticaria, arthralgia, fever, and especially meningitis seen in AIF-PNH are shared by children with cryopyrinopathies or cryopyrin-associated periodic syndrome (reviewed by Neven et al (36)). Cryopyrinopathies are caused by gain-of-function mutation in *NLRP3* that leads to easy activation of NLRP3 inflammasomes in monocytes and autoinflammatory symptoms (36), further suggesting that inflammasomes are activated in monocytes from AIF-PNH patients. It was reported that autoinflammation occurs in patients having mosaicism with *NLRP3*-mutant cells even when the mutant clone size is small (frequency of mutant allele in whole blood cells being 4.3% to 6.5%) (37). This is relevant to the symptoms/clone size relationship in AIF-PNH as discussed above.

PIGU is an essential component of GPI transamidase, forming a protein complex with PIGT, PIGS, PIGK, and GPAA1 (Figure 1A) (38). PIGU-defective cells do not express GPI-APs on their surface (38). The *PIGU* gene is localized at approximately 7.4 Mb centromeric to the myeloid CDR (Figure 2A and 2B), and regions of somatic deletions of 18 and 12 Mb in GPI-AP-defective cells from J1 and G2, respectively, included the entire *PIGU* gene as well as myeloid CDR and *PIGT* gene. The levels of PIGU protein in these cells would be around half of the normal levels. It appears unlikely that the 50% reduction in PIGU has a significant impact on these cells. The levels of PIGT protein in the same cells would be zero or very low because mutations in the remaining *PIGT* gene are a nonsense mutation (E84X) in J1 and a frameshift mutation (frameshift after G254) in G2. For any remaining PIGT protein, half of the normal amounts of PIGU protein would be excessive for making the GPI transamidase complex. However, it is conceivable that, if a similar somatic deletion including *PIGU* and myeloid CDR occurs in a hematopoietic stem cell of an individual who bears a germline *PIGU* loss-of-function mutation, AIF-PNH caused by *PIGU* mutation might occur.

Germline *PIGT* mutations were reported in patients with IGD (Table S2), which is characterized by developmental delay, seizures, hypotonia, and typical facial dysmorphism (11, 39-43). Inflammatory symptoms and intravascular hemolysis were not reported in IGD patients with *PIGT* mutations. They had either partial loss-of-function homozygous mutations (families 1, 4, and 6), or combinations of a partial loss-of-function mutation and a null or nearly null mutation (families 2, 3, 5, and 7). Therefore, cells from the patients with IGD have only partially reduced PIGT activities and express only partially reduced levels of CD59 and DAF/CD55, and may have free GPI only to a small extent. In contrast, both germline and somatic mutants in AIF-PNH were functionally null or nearly null (Table 1). Therefore, the affected cells from AIF-PNH patients lost CD59 and DAF/CD55 severely or completely and had high levels of free GPI. Interestingly, the same mutation c.250G>T (p.E84X) was found in J1 and two Japanese patients with IGD (11, 43) who were not related to each other. AIF-PNH patient J1 and mothers of two IGD patients from families 2 and 7 had the same heterozygous non-sense *PIGT* mutation (Table S2). These mothers were healthy and no inflammatory symptoms were reported for them (11, 43), suggesting that autoinflammation of J1 was not caused by haploinsufficiency but was initiated after the somatic loss of the other *PIGT* copy occurred. In addition, the reported allele frequency of this variant *PIGT* in the East Asian population is 0.0002316, suggesting that 55,000 Japanese may have this variant (44). Although inheritance of the germline *PIGT* mutations in AIF-PNH patients is not formally proven because DNA samples were not available from their families, it is highly likely that J1 inherited the *PIGT* variant from his mother.

FLAER is a fluorescent non-lytic variant of aerolysin (45) and is conveniently used to stain cell-surface GPI-APs and to determine the affected cells in patients with PNH (45, 46). Aerolysin specifically binds to the GPI moiety of some but not all GPI-APs, and requires simultaneous association with N-glycan for high-affinity binding (47–49). Our results with AIF-PNH cells (Figure 3E, F) indicate that FLAER binds to protein-bound GPI but not to free GPI.

## Materials and Methods

### Blood samples and flow cytometry

Peripheral blood samples were obtained from patients J1 (8), G1 (7), G2 and G3 with AIF-PNH, and six patients with PNH after informed consent. Peripheral blood leukocytes (Figure S1B), erythrocytes and reticulocytes were stained for GPI-APs. T5-4E10 mAb (T5 mAb) against free GPI of *Toxoplasma gondii* was a gift from Dr. J. F. Dubremetz (18). T5 mAb recognizes mammalian free GPI bearing N-acetylgalactosamine (GalNAc) side-chain linked to the first mannose (Figure 3A)(19). T5 mAb does not bind to free GPI when galactose (Gal) is attached to GalNAc. Therefore, reactivity of T5 mAb to free GPI is affected by an expression level of Gal transferase that attaches Gal to the GalNAc. Cells were analyzed by a flow cytometer (MACSQuant Analyzer VYB or FACSCalibur) and FlowJo software.

### DNA and RNA analyses

Granulocytes with PNH phenotype were separated from normal granulocytes by cell sorting after staining by FLAER. DNA was analyzed for mutations in genes involved in GPI-AP biosynthesis by target exome sequencing, followed by confirmation by Sanger sequencing (7). DNA was also analyzed by array comparative genomic hybridization for deletion (7). Methylation status of CpG was determined by bisulfite sequencing and a SNuPE assay (13). Total RNA was extracted with the RNeasy Mini Kit (Qiagen) including DNase digestion and DNA cleanup, and reverse transcription was performed with the SuperScript VILO cDNA Synthesis Kit (Invitrogen). Levels of *L3MBTL1*, *SGK2*, *IFT52*, *MYBL2, ABL* and *GAPDH* mRNAs were analyzed by quantitative real-time PCR (Table S1).

### Cell lines

*PIGT*-defective CHO cells and *PIGL*-defective CHO cells were reported previously (11, 50). CRISPR/Cas 9 system was used to generate *PIGT* and *PIGA* knockout (KO) human monocytic THP-1 cells (ATCC) (Table S1 for guide RNA sequences). KO cells were FACS sorted for GPI-AP negative cells. Each knockout cell was rescued by transfection of a corresponding cDNA.

*SLC35A2* gene was knocked out in PIGTKO THP-1 cells by CRISPR/Cas9 system.

### Inflammasome activation and IL1β measurements

Toll-like receptor 2 (TLR2) ligands, Pam_3_CSK4 and Staphylococcus aureus LTA, are from InvivoGen (23, 51). ATP and MSU for activating inflammasomes are from Enzo Life Sciences and InvivoGen (23, 52). Peripheral blood mononuclear cells were stimulated by Pam_3_CSK4 or LTA for 4 hr at 37℃ and after washing by ATP or MSU for 4 hr at 37℃. IL1β ELISA kit (BioLegend) was used to measure IL1β secreted into the supernatants. Polyclonal rabbit anti-IL1β antibody for western blotting was from Cell Signaling Technology. PIGAKO, PIGTKO and wild-type THP-1 cells were differentiated into adherent macrophages in complete RPMI 1640 medium containing 100 ng/ml phorbol 12-myristate 13-acetate (PMA; InvivoGen) for 3 hr, and then with fresh complete medium for overnight (53). For stimulation, medium was replaced with serum free medium with Pam_3_CSK_4_ (200 ng/ml), followed 4hr-later by ATP stimulation for 4 hr (5 mM).

### Stimulation of THP-1-derived macrophages with complement

As a source of complement, whole blood was collected from healthy donors after informed consent, and serum separated, aliquoted and stored at −80°C prior to use. Inactivation of complement was carried out by heating serum at 56°C for 30 min. To prepare acidified serum (AS) that allows activation of the alternative pathway on the cell surface, 21 volumes of serum was mixed with 1 volume of 0.4 M HCl to have pH of approximately 6.7. C6- and C7-depleted sera and purified C6 and C7 proteins were purchased from Complement Technology. Differentiated cells were stimulated with acidified normal serum, or acidified C6-depleted and C7-depleted sera, and those reconstituted with C6 and C7, respectively, at 37°C for 5hr and secreted IL1β was measured by ELISA. C5 was inhibited by addition of 35 μg/ml anti-C5 mAb (eculizumab, Alexion Pharmaceuticals).

For ex vivo blockade of human C5aR, anti-human C5aR or nonpeptide C5aR antagonist W-54011 (5 μM, Merck Millipore)(25) was used. Complement C3 fragments and MAC deposited on the cells were measured by flow cytometry. THP-1 cells were suspended in 20 µl FACS buffer (PBS, 1% BSA, 0.05% sodium azide) with 1:20 human TruStain FcX^TM^ (Fc receptor blocking solution) at room temperature for 10 min. Cells were stained with anti-C3/C3b/iC3b/C3d mAb (clone 1H8, BioLegend) or rabbit anti-human SC5b-9 (MAC) polyclonal antibodies (Complement Technology) in FACS buffer. After washing twice, cells were incubated with the PE-conjugated goat anti-mouse IgG (BioLegend) or Alexa Fluor488-conjugated goat anti-rabbit IgG (Thermo Fisher) secondary antibody. The anti-human SC5b-9 polyclonal antibodies positively stained PMA-differentiated THP-1 cells without incubation in AS. The same antibodies did not stain similarly differentiated PIGTKO and PIGAKO THP-1 cells, suggesting that the antibody product contained antibodies reacted with some GPI-AP expressed on THP-1-derived macrophages (Figure S8B, C). Because of this reactivity to non-MAC antigen(s), the anti-SC5b-9 antibodies were used for PIGTKO and PIGAKO cells but not for WT cells in experiments shown in Figure 5D and F.

### Statistical analyses

All experiments with THP-1 cells were performed at least three times. All values were expressed as the mean ± SD of individual samples. For two-group comparisons between PIGTKO and PIGAKO cells, Student’s *t*-test was used. *P* values below 0.05 were considered statistically significant.

### Study approval

This study was approved by institutional review boards of Osaka University (approval number 681), University of Ulm (approval numbers 279/09 and 188/16) and University of Berlin (approval number EA2/077/12).

### Data Sharing Statement

All data supporting the findings are available from the corresponding authors.

## Acknowledgements

We thank Drs. Morihisa Fujita (Jiangnan University), Tatsutoshi Nakahata (Kyoto University), Hidenori Ohnishi (Gifu University), Tatsuya Saitoh (Tokushima University) and Yusuke Maeda (Osaka University) for discussion, Dr. Jean-Francois Dubremetz (Montpellier University) for T5-4E10 mAb, and Keiko Kinoshita, Kana Miyanagi, Saori Umeshita and Miguel Rodriguez de los Santos for technical help and Dr. med. Lisa A. Gerdes (Munich University) for collaboration regarding patients G3 as well as the patients for providing blood samples and pictures. We thank Edanz (www.edanzediting.co.jp) for editing the English text of a draft of this manuscript. This work was supported by JSPS and MEXT KAKENHI grants (JP16H04753 and JP17H06422) to TK and a grant from the Japan Society of Complement Research to YM.

## Authorship Contributions

BH, YM, NI, HS, PMK, and TK designed research. YM, MO, AK, TH, Shogo M, TE, MJ, RF, and AH performed research. BH, MK, MA, Sho M, YU, and NK acquired the data. BH, YM, MO, MK, JN, YK, NK, HS, and PMK analyzed data. BH, YM, MO, HS, PMK, and TK wrote the paper.

## Conflict-of-interest statements

BH, YU, TK: honoraria (Alexion Pharma); JN: honoraria (Alexion Pharma), research funding (Japan PNH Study Group); YK: research funding (Chugai Pharmaceutical), honoraria (Alexion Pharma); HS: honoraria and research support (all to University of Ulm from Alexion Pharma).

## Supplementary Information

### Supplemental Methods

#### Blood samples and flow cytometry

Total peripheral blood cells were collected by centrifugation and a small sample was saved for staining erythrocytes. Total blood cells were subjected to hypotonic lysis in 40 volumes of ACK buffer to lyse erythrocytes and the remaining total leukocytes were washed with FACS buffer by centrifugation. Peripheral blood leukocytes were stained for GPI-anchored proteins (GPI-APs) by fluorescence-labeled non-lytic aerolysin (FLAER) (Cederlane), anti-CD14 (clone MOP9, BD Biosciences), -CD16 (clone 3G8, BioLegend), -CD24 (clone ML5, BioLegend), -CD48 (BJ40, BioLegend), -CD59 (clone 5H8)(1) or -CD66b (clone 80H3, Beckman Coulter Immunotech). Erythrocytes and reticulocytes were stained by anti-CD58 (clone AICD58, Beckman Coulter) or -CD59 (clone p282/H19, BD). Lymphocytes, monocytes and granulocytes were differentiated based on forward and side scatters (Figure S1B). In the lymphocytes gate, B- and T-lymphocytes were further gated by CD19- or CD3-positivity, respectively. For this, leukocytes were stained with either Pacific Blue-conjugated anti-CD19 or anti-CD3 (clones HIB19 and OKT3, BioLegend) together with staining for free GPI and GPI-APs. Other antibodies used were PE-anti-CD55/DAF (clone IA10, BD), PE-anti-CD88/C5aR (BioLegend) and anti-TfR (clone H68.4, Thermo Fisher). T5-4E10 mAb (T5 mAb) for free GPI (2) is now available from BEI Resources, NIAID, NIH.

For DNA and RNA analyses, granulocytes with PNH phenotype and normal granulocytes were separated by a cell sorter (FACSAria, BD). For this, the total blood leukocytes were stained by FLAER and subjected to cell sorting. Granulocytes were identified by their side and forward scatters and were separated into FLAER-negative and -positive fractions.

#### Determination of break points causing the 18 Mb deletion in patient J1

Genomic DNA of peripheral blood granulocytes from patient J1 containing 29% GPI-AP-deficient cells was used as a template for PCR to amplify a fragment including break points. All combinations of forward primers 1 to 3 and reverse primers 4 to 6 were used (positions and sequences of PCR primers in Figure S5A, top and Table S1). A fragment amplified with primers 1 and 5 (red arrowheads) was Sanger sequenced using primers 7 to 11 (positions and sequences in Figure S5A, top and Table S1).

#### Quantitative PCR (qPCR) analysis of chromosome 20q bearing the break points to determine the clone size of GPI-AP-deficient cells in patient J1

Two samples of J1’s genomic DNA, one (sample #1) prepared from the whole blood leukocytes taken four months before the onset of recurrent hemolysis and the other (sample #2) prepared from granulocytes (29% GPI-AP-deficient) taken one month after the commencement of eculizumab therapy, were analyzed by qPCR. Primers to amplify a region including the break were designed based on the determined break points (Figure S5A). A region in exon 3 of PIGL was amplified as a reference. Primer sequences are shown in Table S1. SYBR Green Master Mix and StepOnePlus system (Thermo Fisher Scientific) were used. Ratios of (Break)/(PIGL) were determined for #1 and #2 samples with the 2(−ΔΔCt) method for comparative analysis, and a relative level of the break in #1 sample was determined setting a ratio of #2 sample as 1 (Results are shown in Figure 3G). Percent GPI-AP-deficient cells in the whole blood leukocytes at four months before the onset of recurrent hemolysis was determined by multiplying 29% by a relative level of the break in #1.

#### Generation and characterization of PIGTKO, PIGAKO and PIGT-SLC35A2 double KO THP-1 cells.

Gene disruption in THP-1 cells was made with CRISPR/Cas9 system (guide RNA sequences shown in Table S1). PIGTKO and PIGAKO cell clones were established based on their phenotypes (loss of FLAER- and CD55-positive staining) and mutations as assessed by Sanger sequencing. Specificities of KO were confirmed by restoration of the wild-type phenotypes after transfection of corresponding cDNAs. PIGT-SLC35A2 double KO (PIGT-SLC35A2KO) cells were established by KO SLC35A2 in PIGTKO cells and sorting T5 mAb staining positive cells.

#### Quantitative RT-PCR (qRT-PCR) for IL1β and NLRP3

Levels of IL1β and NLRP3 mRNAs were analyzed by qRT-PCR using SYBR Green (Applied Biosystems) and appropriate primers (Table S1). Data were normalized to β–actin according to the 2(−ΔΔCt) method for comparative analysis. IL1β and NLRP3 mRNA levels of PMA-treated WT cells were set to 1. Result was expressed as the mean ± SD of duplicate determinations of a representative experiment.

### Supplemental Results

#### Characterization of the patients’ blood samples and bone marrow

Blood cell counts and neutrophil percentage in leukocytes of patients G1 and G3 were shown as a function of time in days in Figure S1A. LDH levels of patient G1 were also shown. Total leukocyte counts and platelet counts were within normal ranges in patients G1 and G3 (Figure S1A) and J1 (3), except at around severe hemolysis. The bone marrow specimen of patient G3, collected 6 months after the first meningitis episode and 4 years before the diagnosis of PNH showed an increased hematopoiesis. There were no signs of malignancy or aplasia. Taken together with reports that there were no signs of aplasia in the bone marrow of J1 (3) and G1 (4), bone marrow failure does not seem to be a feature of patients with autoinflammation-PNH (AIF-PNH) characterized by PIGT mutations.

#### Determination of break points causing the 18 Mb deletion in patient J1

The 18 Mb deletion was previously determined by a SNP array analysis of GPI-AP-deficient and GPI-AP-positive granulocytes from J1 (3). The SNP array analysis showed that SNPs at positions 31,905,813 and 49,909373 were present in the deletion-containing chromosome 20 of GPI-AP-deficient cells whereas all SNPs between those at 31,925,918 and 49,759,935 were absent (Figure S5A, top : positions according to GRCh37/hg19, NCBI/UCSC). Therefore, the centromeric and the telomeric break points must exit between positions 31,905,813 and 31,925,918, and positions 49,759,935 and 49,909373, respectively. To amplify a DNA fragment including the break points, we designed three PCR primers each in the centromeric (primers 1 to 3) and the telomeric (primers 4 to 6) regions (Figure S5A, top). Using genomic DNA from J1 granulocytes containing 29% GPI-AP-deficient cells as a template (sample #2 in Figure S5B, right part), PCR with primers 1 and 5 (primer set 1-5) generated a 6-kbp product (marked *) whereas PCRs with all other primer sets gave negative results or non-specific products. To determine nucleotide sequence of the 6-kbp PCR product, we generated five sequence primers (7 to 11 in Figure S5A, top). Sequencing with primer 8 (red) was successful. Two break points at positions 31,909,463 and 49,798,761, and an insertion of five nucleotides (5’-ACATT) between the break points were identified (Figure S5A, bottom).

#### A clone size of GPI-AP-negative leukocytes in patient J1 at four months before the onset of recurrent hemolysis

We also used J1’s genomic DNA prepared from the whole blood leukocytes taken four months before the onset of recurrent hemolysis (sample #1 in Figure S5B, left part) as a template in PCR. Similar to sample #2, primer set 1-5 generated a 6-kbp product (marked *) whereas all other primer sets gave negative results or non-specific products (Figure S5B, left part). The 6-kbp product amplified from sample #1 was confirmed to be the same break-containing region by Sanger sequencing. The amount of the 6-kbp product from sample #1 was much smaller than that from sample #2 (29% GPI-AP-deficient cells). These results indicate that J1 had a small GPI-AP-clone four months before the onset of PNH.

To determine the clone size at four months before the onset of recurrent hemolysis, we analyzed samples #1 and #2 by qPCR. The level of the break-containing DNA in sample #1 was approximately 10% of sample #2 (Figure 3G), indicating approximately 3% of total leukocytes were GPI-AP-deficient.

#### Characterization of PIGTKO, PIGAKO and PIGT-SLC35A2 KO THP-1 cells

The surface expressions of GPI-APs were lost from both PIGTKO and PIGAKO cells and were restored by transfection of PIGT and PIGA cDNAs, respectively (Figure S6A). PIGTKO, PIGAKO and wild-type (WT) THP-1 cells were differentiated into macrophages by phorbol 12-myristate 13-acetate (PMA) to induce competence in inflammasome activation (5). They were then tested by priming with Pam_3_CSK_4_ followed by 4h-treatment with ATP. IL1β was released at similar levels from WT, PIGTKO and PIGAKO cells (167.9±61.5, 144.3±19.7, 200.3±48.8 pg/ml), indicating that they have comparable abilities to respond to inflammasome activators (Figure S6B). PIGT-SLC35A2KO cells were strongly stained by T5 mAb whereas PIGTKO cells were only very weakly stained by T5 mAb (Figure S6C). This result indicated that most free GPIs in PIGTKO cells were not detected by T5 mAb because of capping by galactose (schematic shown in Figure 3A).

#### IL1β and NLRP3 levels in PIGTKO, PIGAKO and WT THP-1 cells before and after incubation with complement

Transcriptional induction of IL1β and NLRP3 were at similar levels in PIGTKO, PIGAKO and wild-type (WT) THP-1 cells and the PIGTKO cells rescued by PIGT cDNA. IL1β was induced by PMA alone, while NLRP3 induction was further enhanced by the acidified serum (Figure S7A).

**Fig. S1.**
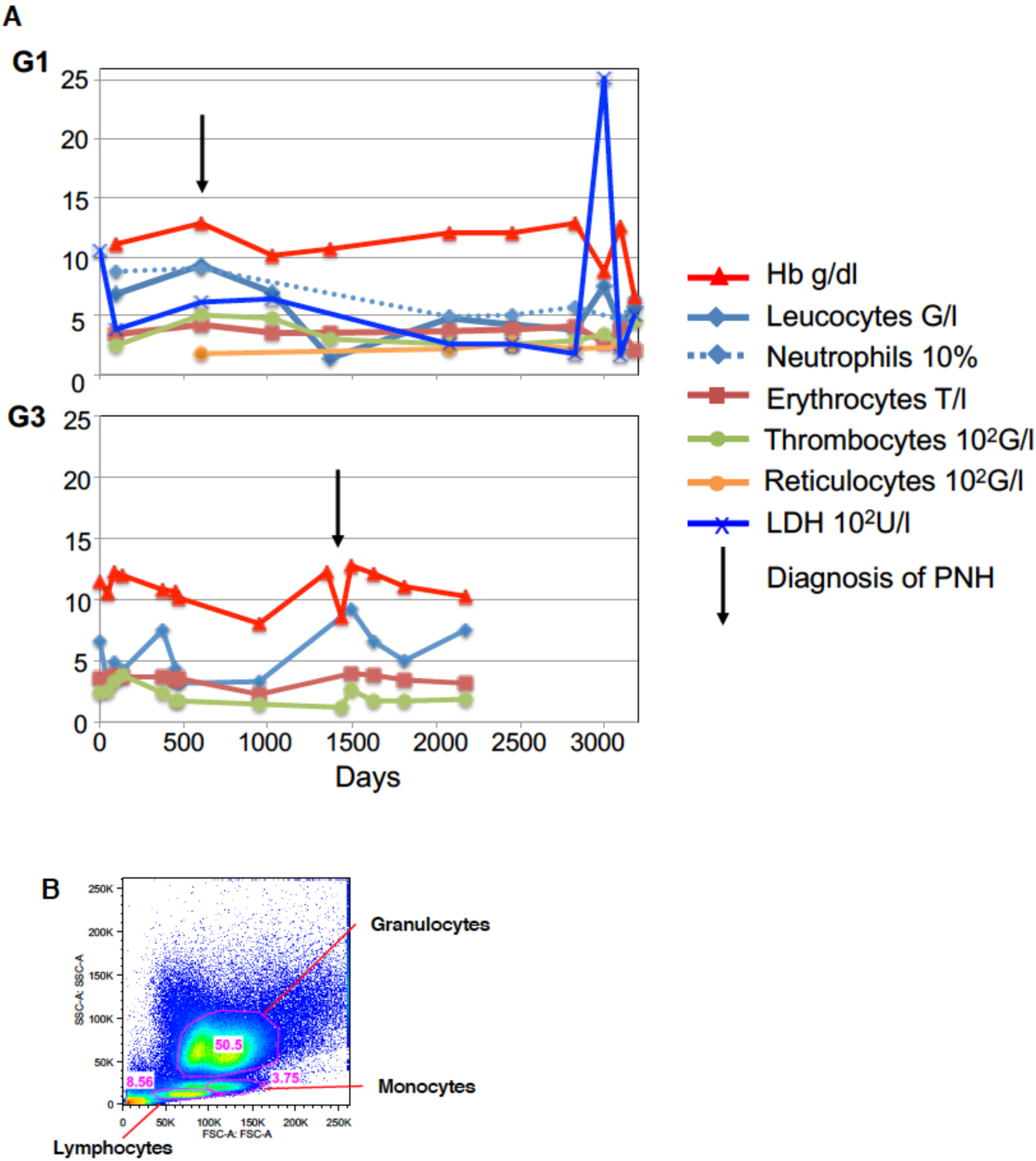
Blood cell counts in patients G1 and G3 and gating of peripheral blood cells. **A.** Time course of blood cell counts. A very high peak of LDH found in patient G1 on around day 3000 was caused by severe hemolytic crisis occurred after a transient extension of the eculizumab therapy interval (4). G/l, billion cells per liter; 10^2^G/l, hundred billion cells per liter; T/l, trillion cells per liter. **B.** Gating of granulocyte, monocyte and lymphocyte populations based on forward (FSC) and side (SSC) scatters.

**Fig. S2.**
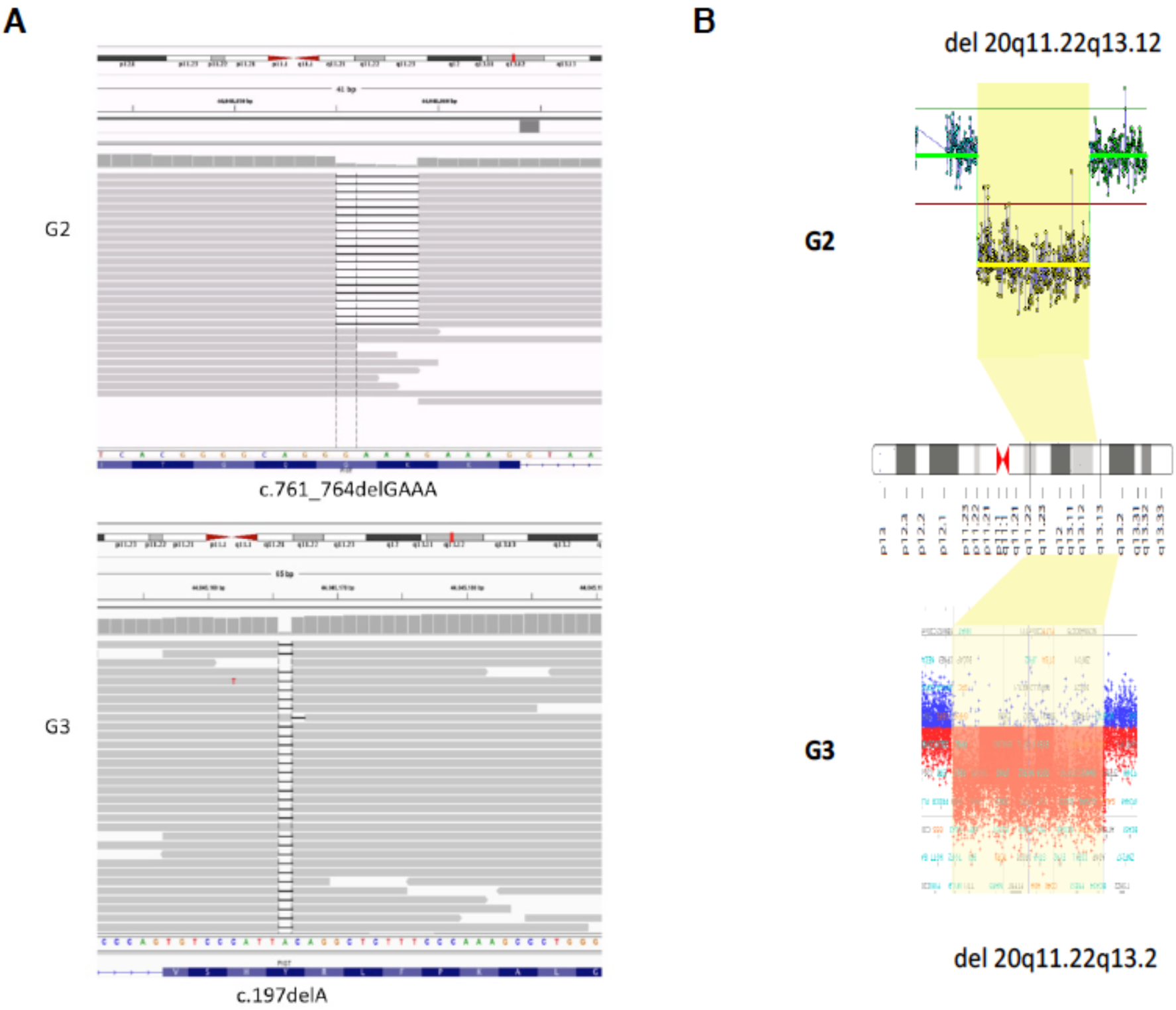
Rare pathogenic variants and somatic mutations of *PIGT* in AIF-PNH G2 and G3 revealed by Gene Panel analysis and array comparative genomic hybridization. A. The inherited variants in G2 and G3 are a four-base pair deletion (NM_015937: c.761_764delGAAA) and a one-base pair deletion (NM_015937: c.197delA) in *PIGT*. **B.** The somatic deletion in G2 spans over 12 Mb on chr 20q and involves *PIGT* and *PIGU*, arr[hg19] 20q11.22q13.12(32,837,289-44,973,145)x1. The somatic event in G3 is a 15 Mb deletion involving *PIGT*, arr[hg19] 20q11.22q13.2(34,975,105-50,140,767) x1.

**Figure S3.**
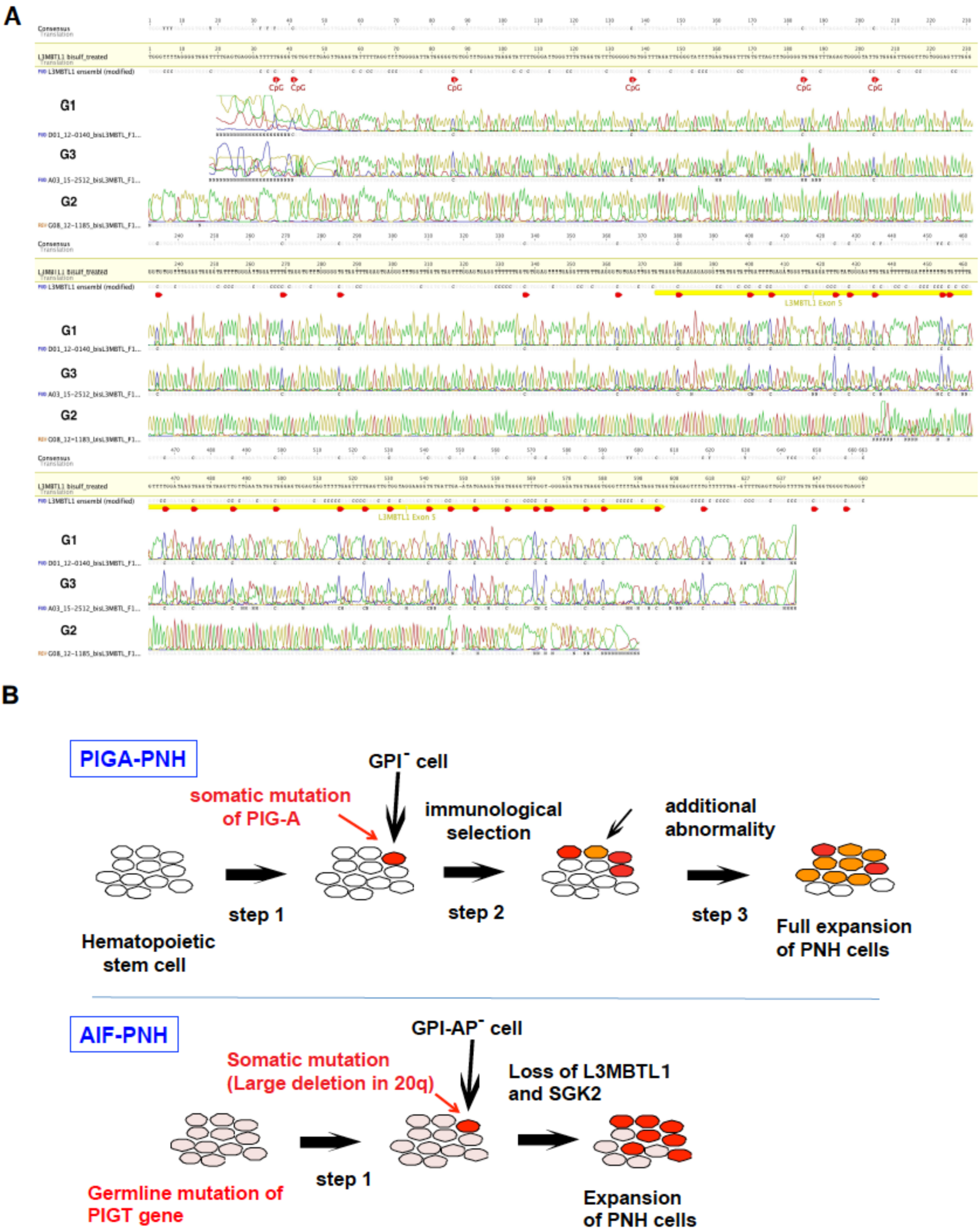
Methylation status of *L3MBTL1* and a model for clonal expansion in AIF-PNH. **A.** Bisulfite sequencing of the informative region in *L3MBTL1* in G1, G2 and G3. **B.** Models of clonal expansion of GPI-defective cells in PNH (top) and AIF-PNH (bottom). In PNH, somatic mutation of *PIGA* gene in a hematopoietic stem cell generates GPI-defective clone (step 1). Under bone marrow failure conditions, normal stem cells are damaged by autoimmune mechanisms whereas GPI-defective clone survives, causing an initial clonal expansion (step 2). The GPI-defective subclone that acquired benign-tumor-like growth phenotype expands greatly (step 3). In AIF-PNH, patients had a germ line loss-of-function mutation in the maternal allele of *PIGT*. A deletion spanning the entire *PIGT* and myeloid common deletion region occurs in the paternal allele in a hematopoietic stem cell, generating GPI-AP-defective clone (step 1). Because of losses of maternally imprinted *L3MBL1* and *SGK2* genes, the GPI-AP-defective clone may obtain a competence to expand and may initially contribute to non-erythrocytic myeloid cells causing recurrent autoinflammation. Years later, *PIGT*-defective clone may begin to generate sufficient numbers of GPI-AP-defective erythrocytes to cause PNH phenotype.

**Figure S4.**
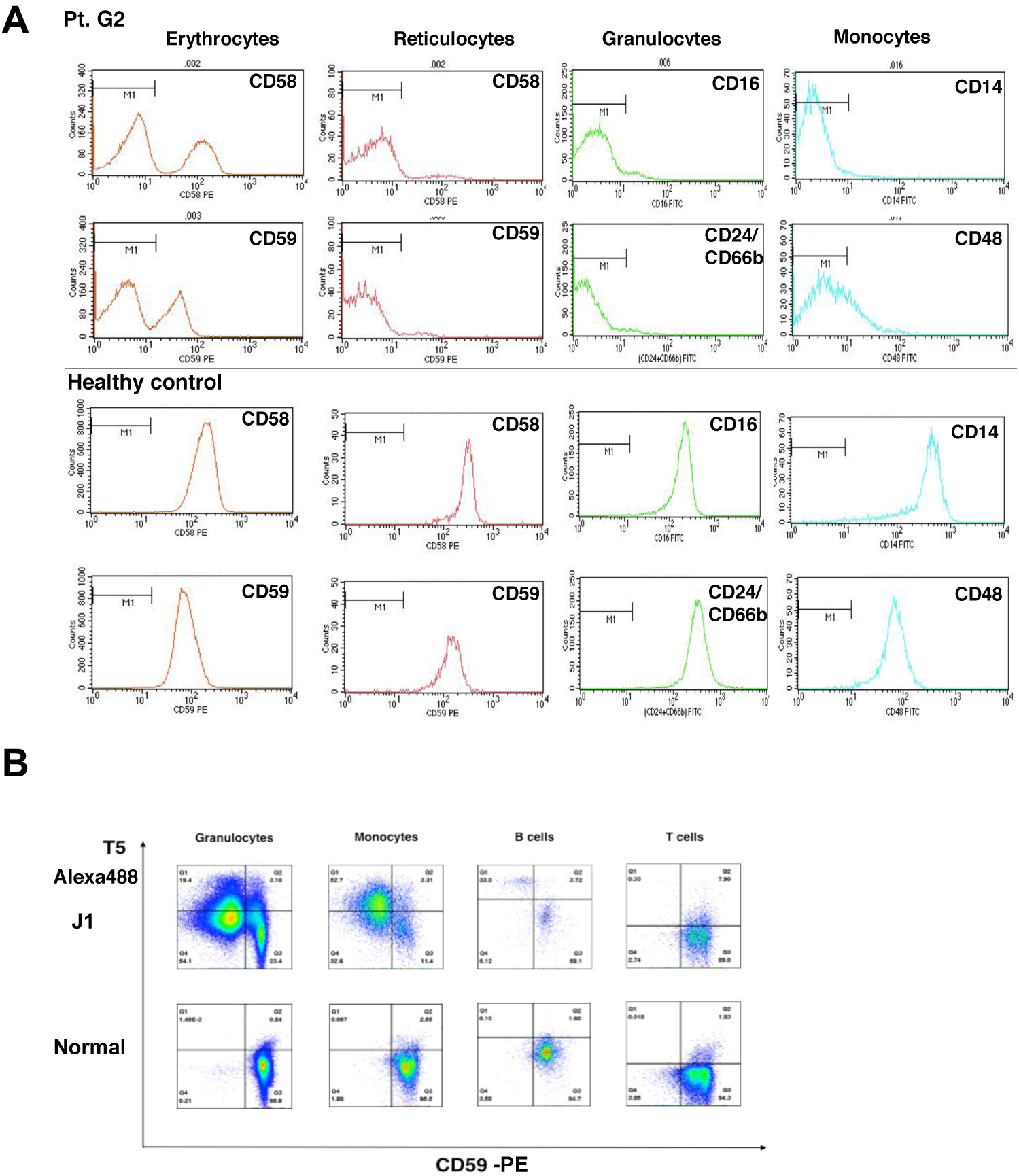
Flow cytometry of blood cells from patients with AIF-PNH. **A.** Peripheral blood cells from G2 with AIF-PNH (top two lows) and a healthy control (bottom two lows) were stained for GPI-APs. **B.** Granulocytes, monocytes, B- and T-lymphocytes from J1 with AIF-PNH and a normal individual, stained by T5 mAb and anti-CD59 mAb.

**Figure S5.**
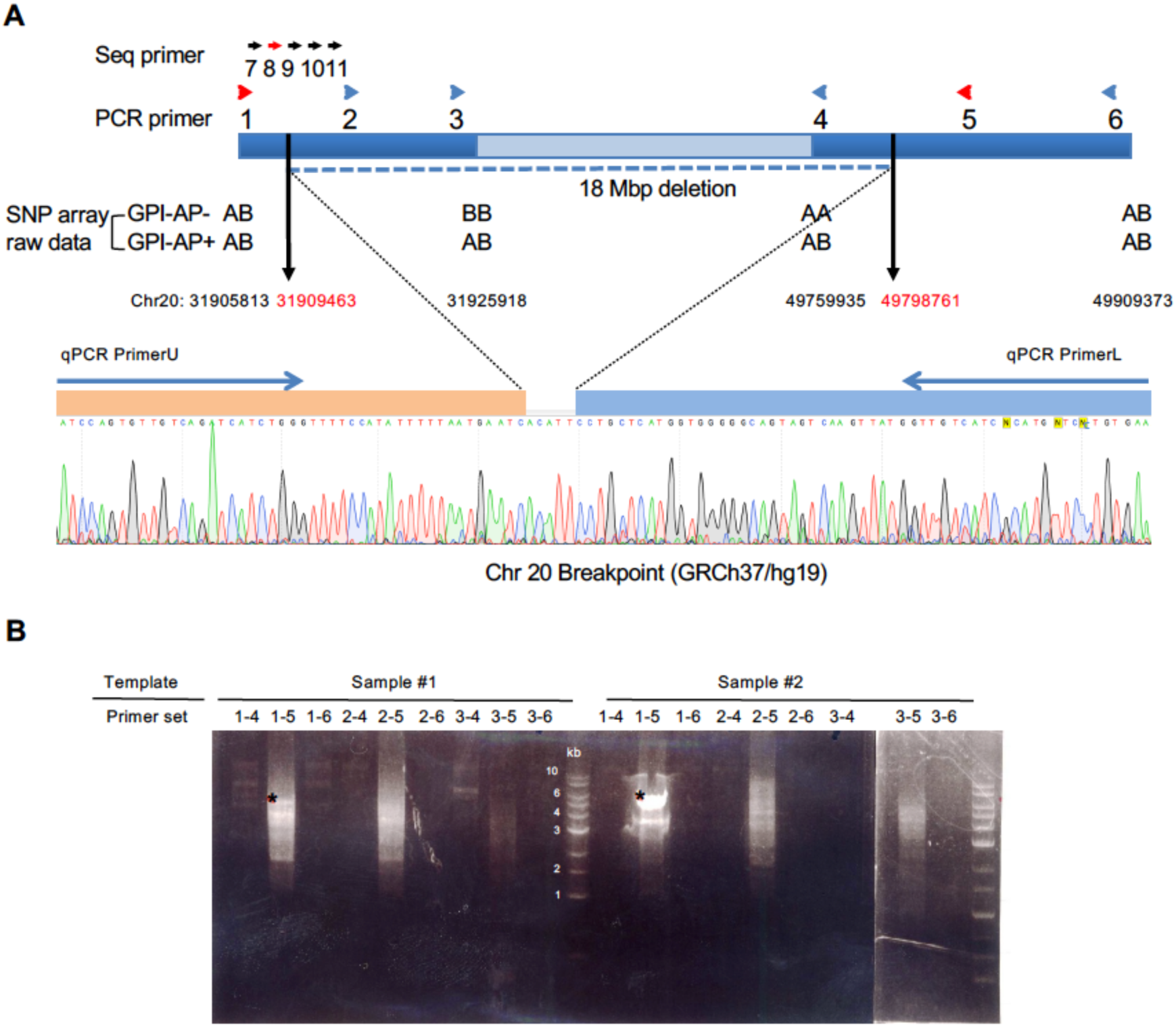
Determination of the break points causing the 18 Mb deletion in J1. **A** (Top) Schematic of a region in patient J1’s chromosome 20q bearing the 18 Mb deletion (centromere on the left). Four informative SNPs (AB, heterozygous; AA and BB, homozygous) are shown in their position numbers below the schematic: SNPs heterozygous in both GPI-AP-positive and GPI-AP-deficient cells (at 31,905,813 and 49,909,373) were present in GPI-AP-deficient cells whereas those homozygous in GPI-AP-deficient cells and heterozygous in GPI-AP-positive cells (at 31,925,918 and 49,759,935) were within the deleted region in GPI-AP-deficient cells. PCR primers 1 to 6 used to amplify a region including break points are shown immediately above the schematic. Sequence primers 7 to 11 used to determine nucleotide sequence of the PCR product with primers 1 and 5 (red arrowheads) (see **B** for PCR products) are shown on top: sequencing with primer 8 (red) was successful. (Bottom) Sanger sequence result with primer 8. Two break points at positions 31,909,463 and 49,798,761, and an insertion of five nucleotides (5’-ACATT) between the break points were identified. Locations of primers for qPCR (Primers U and L) to determine the clone size of GPI-AP-cells are shown above the sequence. **B.** PCR products from template samples #1 and #2 with all combinations of primers. Sample #1, genomic DNA from the whole blood leukocytes taken four months before the onset of recurrent hemolysis; sample #2, genomic DNA from granulocytes (29% GPI-AP-deficient) taken one month after the commencement of eculizumab therapy. PCR with primer set 1-5 generated a 6 kbp product strongly from sample #2 and weakly from sample #1. All other primer sets gave negative results or non-specific products.

**Figure S6.**
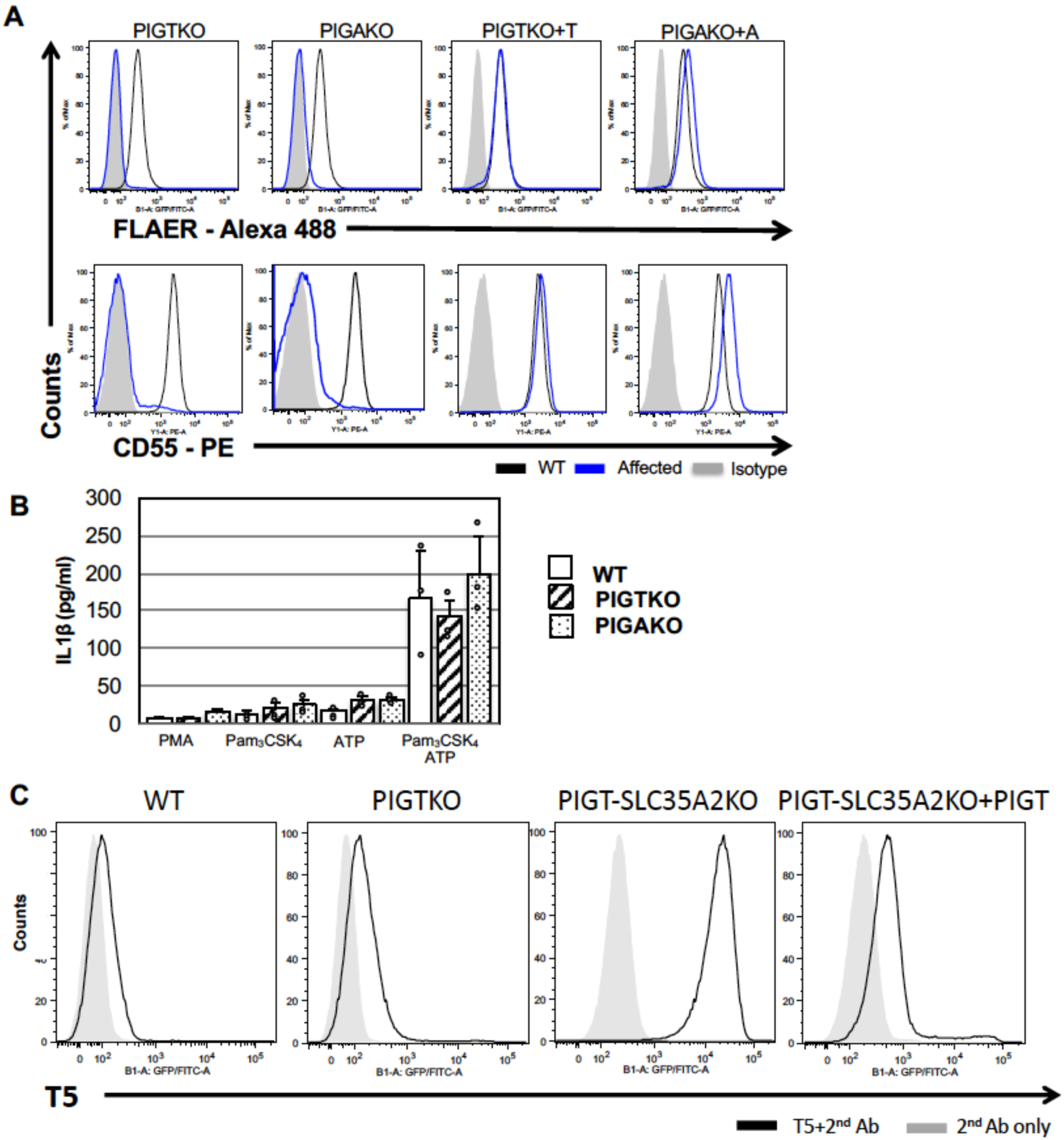
Characterization of PIGT and PIGA knockout THP-1 cells. **A.** Surface expression of GPI-APs on PIGT knockout (KO), PIGAKO and PIGT-/ PIGA-rescued (PIGTKO+T, PIGAKO+A) THP-1 cells. Test cells (blue lines) and wild-type cells (solid lines) were stained with FLAER (top panels) and PE-labeled anti-CD55 antibody (bottom panels). Negative staining controls (shaded) for FLAER and anti-CD55 were buffer only and isotype matched monoclonal antibody, respectively. The expressions of GPI-APs on PIGTKO and PIGAKO cells were lost and normalized by rescue transfections of relevant cDNAs. **B.** Secretion of IL1β from THP-1 cells. WT, PIGTKO and PIGAKO THP-1 cells were differentiated into macrophages by PMA (100 ng/ml). The differentiated cells were primed with Pam_3_CSK_4_ (200 ng/ml) for 4 hr, after which medium was removed and pulsed for 4 hr with ATP (5mM). IL1β released into medium was measured by ELISA. Data are expressed as mean + SD of three independent experiments. **C.** Expression of free GPI on WT, PIGTKO, PIGT-SLC35A2 KO, PIGT-rescued PIGT-SLC35A2 KO THP-1 cells. T5 mAb did not positively stain WT and PIGTKO THP-1 cells whereas it stained PIGTKO-SLC35A2KO THP-1 cells strongly. SLC35A2KO cells derived from PIGTKO cells are defective in galactosylation because of defective UDP-Gal transporter encoded by *SLC35A2*, indicating that PIGTKO THP-1 cells express free GPI whose GalNAc side-chain is fully galactosylated.

**Figure S7.**
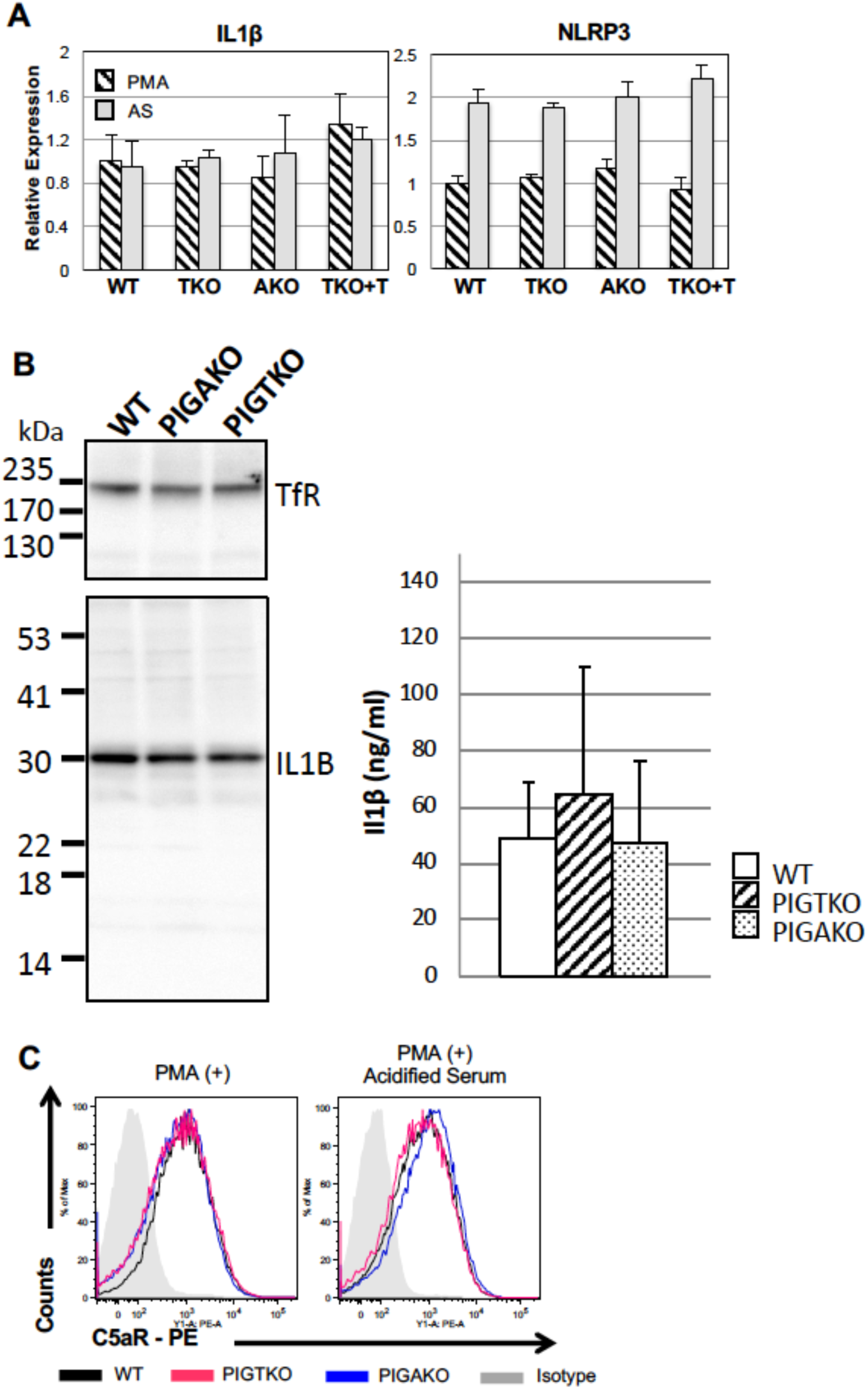
Responses of THP-1 cells to complement. **A.** Levels of IL1β and NLRP3 transcripts in WT, PIGTKO, PIGAKO and PIGT-rescued PIGTKO THP-1 cells. Cells were differentiated to macrophages by PMA and RNA prepared before (hatched bars) and after (grey bars) 5-hr incubation in 50% acidified serum (AS). IL1β (left) and NLRP3 (right) transcripts were measured by qRT-PCR. Data are expressed as mean + SD of triplicate samples from one of three independent experiments. **B.** Levels of IL1β protein in WT, PIGTKO and PIGAKO THP-1 cells. After differentiation by PMA, detergent extracts were prepared and IL1β determined by western blotting (left) and ELISA (right). TfR, transferrin receptor used as a loading control. Levels of IL1β determined by ELISA were expressed as mean + SD of three independent experiments. **C.** Levels of C5aR on THP-1-derived macrophages before and after incubation in acidified serum. WT (black lines), PIGTKO (red lines) and PIGAKO (blue lines) cells before (left) and after 5-hr incubation in 50% acidified serum (right) were stained by anti-CD88 mAb. Shaded, WT cells stained by isotype control antibody. Mean + SD of three independent experiments.

**Figure S8.**
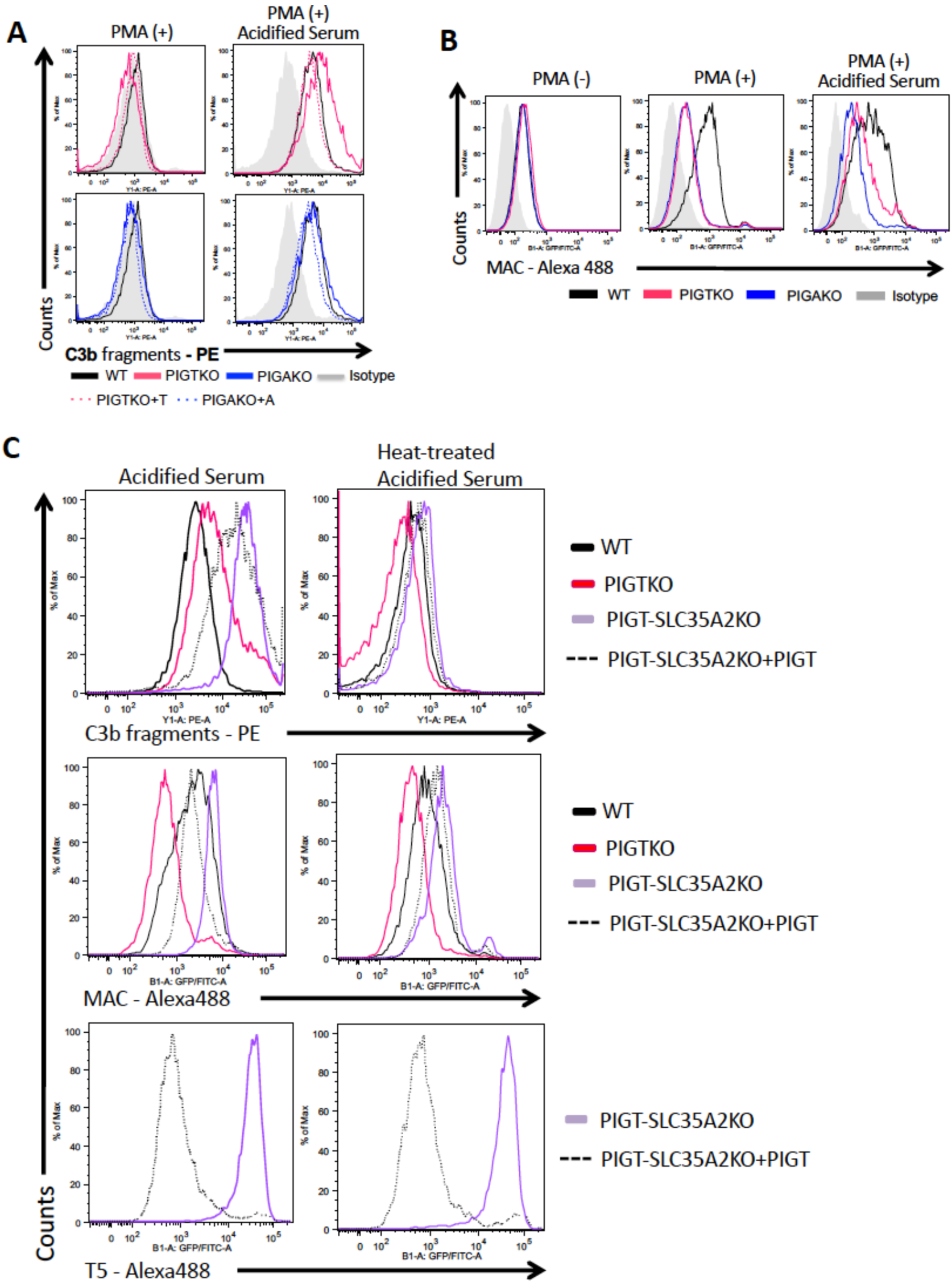
**A.** Detection of C3b fragments by flow cytometry. THP-1 cells were differentiated into macrophages by PMA (left) and incubated with 50% acidified serum for 5 hr after differentiation (right). WT (black lines), PIGTKO (solid red lines), PIGT-rescued PIGTKO (dotted red lines), PIGAKO (solid blue lines) and PIGA-rescued PIGAKO (dotted blue lines). One representative experiment of three independent experiments. **B.** Detection of MAC by flow cytometry. WT (black lines), PIGTKO (red lines), PIGAKO (blue lines) THP-1 cells were left undifferentiated (left), differentiated into macrophages by PMA (center) and incubated with 50% acidified serum for 5 hr after differentiation (right). One representative experiment of three independent experiments. **C.** Detection of C3b fragments (top), MAC (middle) and free GPI (bottom) on PIGT-SLC35A2 KO THP-1 cells and PIGT-rescued PIGT-SLC35A2 KO THP-1 cells after PMA-treatment and incubation with acidified serum (left) or heated acidified serum (right). Black lines, WT; red lines, PIGTKO; purple lines, PIGT-SLC35A2 KO; dotted lines, PIGT-rescued PIGT-SLC35A2 KO.

**Table S1.**
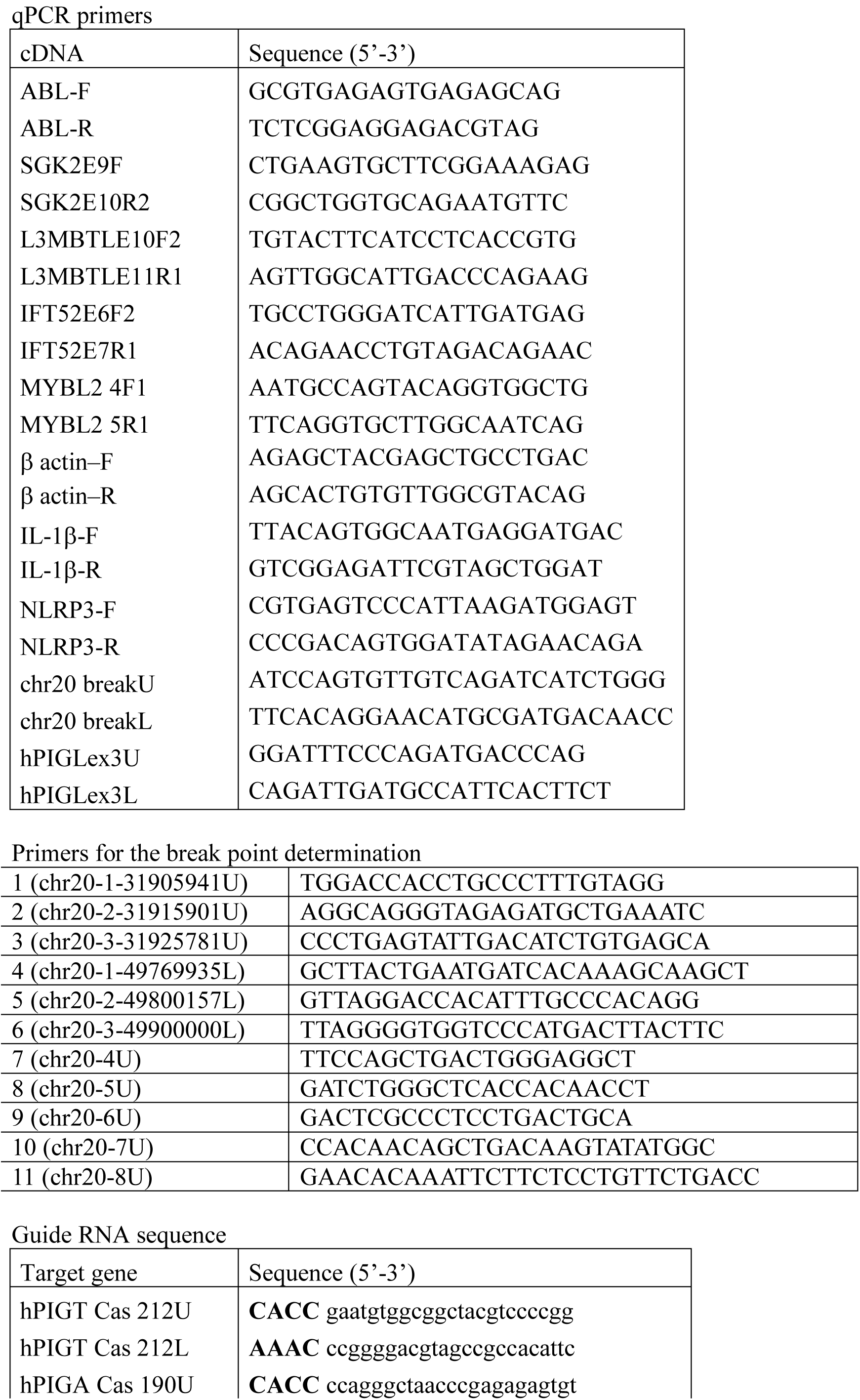
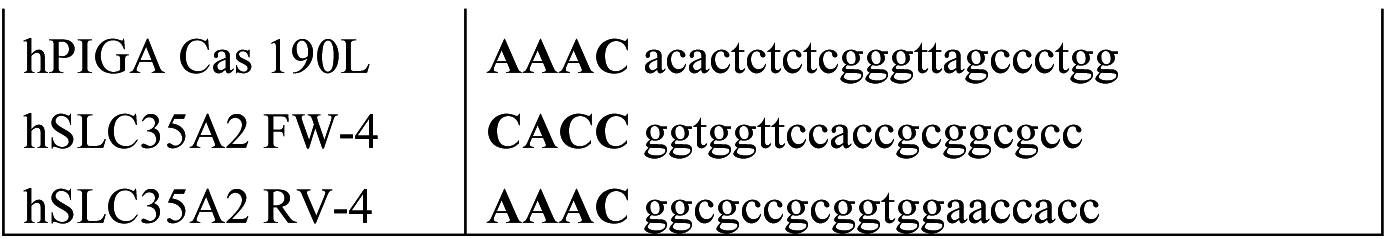
Table S1. Sequences of primers and guide RNA used in this study.

**Table S2.** PIGT mutations in Inherited GPI Deficiency (IGD)

| Family | Origin | M<sup>1)</sup> allele | P<sup>1)</sup> allele | Reference |
| --- | --- | --- | --- | --- |
| 1 | Aramaic | c.547A>C<sup>2)</sup><br>(p.T183P) | c.547A>C<br>(p.T183P) | Kvarnung (6) |
| 2 | Japanese | c.250G>T<sup>3)</sup><br>(p.E84*) | c.1342C>T<br>(p.R448W) | Nakashima (7) |
| 3 | Caucasian<br>(M)/African<br>American | c.918dupC<br>(p.V307Rfs*13) | c.1342C>T<br>(p.R448W) | Lam (8) |
| 4 | Somalian | c.1079G>T<br>(p.G360V) | c.1079G>T<br>(p.G360V) | Skauli (9) |
| 5 | Caucasian | c.1582G>A<br>(p.V528M) | c.1730dupC<br>(p.L578fs*35) | Pagnamenta (10) |
| 6 | Afganistani | c.709G>C<br>(p.E237Q) | c.709G>C<br>(p.E237Q) | Pagnamenta (10) |
| 7 | Japanese | c.250G>T<sup>3)</sup><br>(p.E84*) | c.1096G>T<br>(p.G366W) | Kohashi (11) |

<sup>1)</sup> M, maternal; P, paternal
<sup>2)</sup> Based on NM\_015937
<sup>3)</sup> The same mutation was found in patient J1 with autoinflammation-paroxysmal nocturnal hemoglobinuria (AIF-PNH) (Table 1).

